# SuperFi-Cas9 exhibits extremely high fidelity but reduced activity in mammalian cells

**DOI:** 10.1101/2022.05.27.493683

**Authors:** Péter István Kulcsár, András Tálas, Zoltán Ligeti, Sarah Laura Krausz, Ervin Welker

## Abstract

Several advances have been made to SpCas9, the most widely used CRISPR/Cas genome editing tool, to reduce its unwanted off-target effects. The most promising approach is the development of increased fidelity nuclease (IFN) variants of SpCas9, however, their fidelity has increased at the cost of reduced activity. SuperFi-Cas9 has been developed recently, and it has been described as a next-generation high fidelity SpCas9 variant, free from the drawbacks of the first-generation IFNs. In this study, we characterized the on-target activity and the off-target propensity of SuperFi-Cas9 in mammalian cells comparing it to first-generation IFNs. SuperFi-Cas9 demonstrated strongly reduced activity but exceptionally high fidelity exhibiting features that are in many aspects similar to those of the first-generation variants, such as evo- and HeFSpCas9. When combined with ABE8e, SuperFi-Cas9 produced DNA editing with high activity rate as well as high specificity by reducing both bystander and SpCas9-dependent off-target base editing.

## Introduction

Several approaches have been developed^1^ to reduce the off-target effects^2^ of the SpCas9 nuclease of the type II CRISPR system, which is the most commonly used tool for gene editing applications. Besides approaches that increase the length of the recognition sequence of the nuclease^3–6^ or ones that limit its activity in space and/or time^7–10^, mutations introduced to the sequence of the ribonucleoprotein complex, affecting either the RNA or the protein component, also seem to be an effective way to increase the specificity of editing. Alteration of the spacer, such as truncation^11^ or extension by one^12^ or two^13^ 5’ G nucleotides as well as the incorporation of modified bases into its sequence, increases the fidelity of SpCas9^14^. Mutations weakening the interactions between the protein and either the targeted^15^ or nontargeted DNA strand^16^, or the spacer^12^, as well as ones weakening the intermolecular interactions between the domains of the protein^17^ lead to higher fidelity. Certain amino acid mutations of SpCas9, identified in selection schemes^18–20^, also result in improved discrimination between on-target and off-target sequences. It has been shown that variants with the above-mentioned types of protein mutations become more selective of targets they are active on in exchange for higher fidelity; the higher the fidelity, the more selective the variant is with targets it is active on^12,21–23^. Interestingly, this target-selectivity translates into either fully or partially reduced activity at some targets, while at other targets these increased-fidelity nucleases (IFNs) show wild-type-(WT)-like activity^12,15,17,18,21,22^. Ultimately, their overall average on-target activity is reduced.

A recent paper has introduced a newly developed variant that can go beyond this paradigm^24^. The variant was generated by rational design exploiting a cryo-electron microscopy structure of SpCas9 with a PAM-distal mismatching single guide RNA (sgRNA). The structure revealed that a flexible loop of the RuvC domain stabilizes the distorted end of the target DNA-sgRNA hybrid helix allowing SpCas9 activation even with mismatches. The authors speculated that the residues, stabilizing the distorted helix end, do not participate in interactions in any known SpCas9 structure complexed with on-target DNA. They proposed that by disrupting these mismatch-stabilizing interactions, off-target cleavage activity of SpCas9 can be diminished without affecting on-target cleavage. Based on this, a new increased-fidelity SpCas9 variant was developed by mutating the seven contacting residues to aspartic acid. It was found to exhibit WT-like on-target activity and decreased off-target activity *in vitro* using a target/off-target pair on which the activity of two IFNs, SpCas9-HF1 and Hypa-SpCas9, had previously been reported to be two orders of magnitude lower than the WT. The authors named this next-generation variant SuperFi-Cas9, inspired by its both high fidelity and on-target activity. Based on this rationale, further next-generation IFNs may be generated with features distinct to those of the first-generation IFNs that exist to date.

As reported earlier, every IFN exhibits a variety of on-target activity rates in a target-dependent manner, ranging from WT-like on-target activity to close to zero activity^12,15,17,18,21,22^. Whether SuperFi would demonstrate uncompromised on-target activity on other targets too is yet unknown. Furthermore, since some of the most important applications of SpCas9 variants exploit its activity in mammalian cells, the most important question raised by the study of Bravo et al. is whether SuperFi will display the same features in mammalian cells. We also speculate that the rationale by which SuperFi was generated implies that its mutations will cause reduced activity on off-target sequences where the mismatches are located at positions 17-20, but on sequences with mismatches existing only at other PAM-distal positions, SuperFi will likely show WT-like activity. To answer these questions, we characterized SuperFi for on-target and off-target activity in mammalian cells, comparing it to appropriate first-generation IFNs (Supplementary Fig. 1a). These experiments showed that SuperFi possesses extremely high fidelity and a reduced activity level that is typical of the highest fidelity first generation IFNs, such as evo- and HeFSpCas9. SuperFi is routinely applicable with 21G-sgRNAs, demonstrates high base editing activity with ABE8e and is capable of mitigating its bystander and off-target activities.

## Results

The on-target activity of SuperFi was examined in an EGFP disruption assay on 25 (Fig. 1a and Supplementary Fig. 1b) and by amplicon sequencing on 20 genomic targets (Fig. 1b and Supplementary Fig. 1c). SuperFi showed significantly lower average on-target activity than WT, Hypa- or SpCas9-HF1, although, at very few targets its activity approached that of the WT nuclease. Western blot (Supplementary Fig. 1d) and expression plasmid titration analyses (Supplementary Fig. 1e) showed that this lower activity could not be explained by lower expression levels. To assess its fidelity, we selected challenging targets^12,15,17,18,25^ that many nuclease variants have failed to edit without off-target activity, as well as targets that have off-targets with mismatches exclusively at positions 18-20 or 14-17 according to earlier GUIDE-seq experiments^12,25^. SuperFi showed superior fidelity on many of these targets in comparison to Hypa- and SpCas9-HF1 (Fig. 1c, d and Supplementary Fig. 2a, b). However, contrary to our expectations we saw increased specificity not only at the 18-20 positions but at the 14-17 positions. The genome-wide off-target effects of SuperFi were examined by GUIDE-seq (Fig. 1e, f). Due to its higher fidelity, instead of Hypa- and SpCas9-HF1, we examined SuperFi in comparison with Blackjack-HypaR- and evoSpCas9 that have more similar fidelity to SuperFi^12^. SuperFi exhibited higher specificity than Blackjack-HypaR- and evoSpCas9 on 3 and 2 out of the 4 examined targets, respectively (Fig. 1e, f and Supplementary Fig. 3). 5’ extended or truncated spacers are frequently applied with SpCas9 nucleases, as these can increase the fidelity of editing^6,11–13,26^, and a 5’ G extension of the spacer (21G-sgRNA) is often necessary to comply with the sequence preference of the U6 and T7 promoters often used for the expression of sgRNAs^12,19^. Only Sniper-Cas9^19^, the lowest fidelity IFN^12^, can be routinely used with 5’-truncated sgRNAs, and only Sniper and the Blackjack variants can be routinely used with 21G-sgRNAs amongst the first-generation IFNs^12,19^. Although truncating the sgRNAs diminished the activity of SuperFi (Supplementary Fig. 4a), it seems to be fully compatible with the extended 21G-sgRNAs (Fig. 1g, h and Supplementary Fig. 4b).

**Figure 1.**
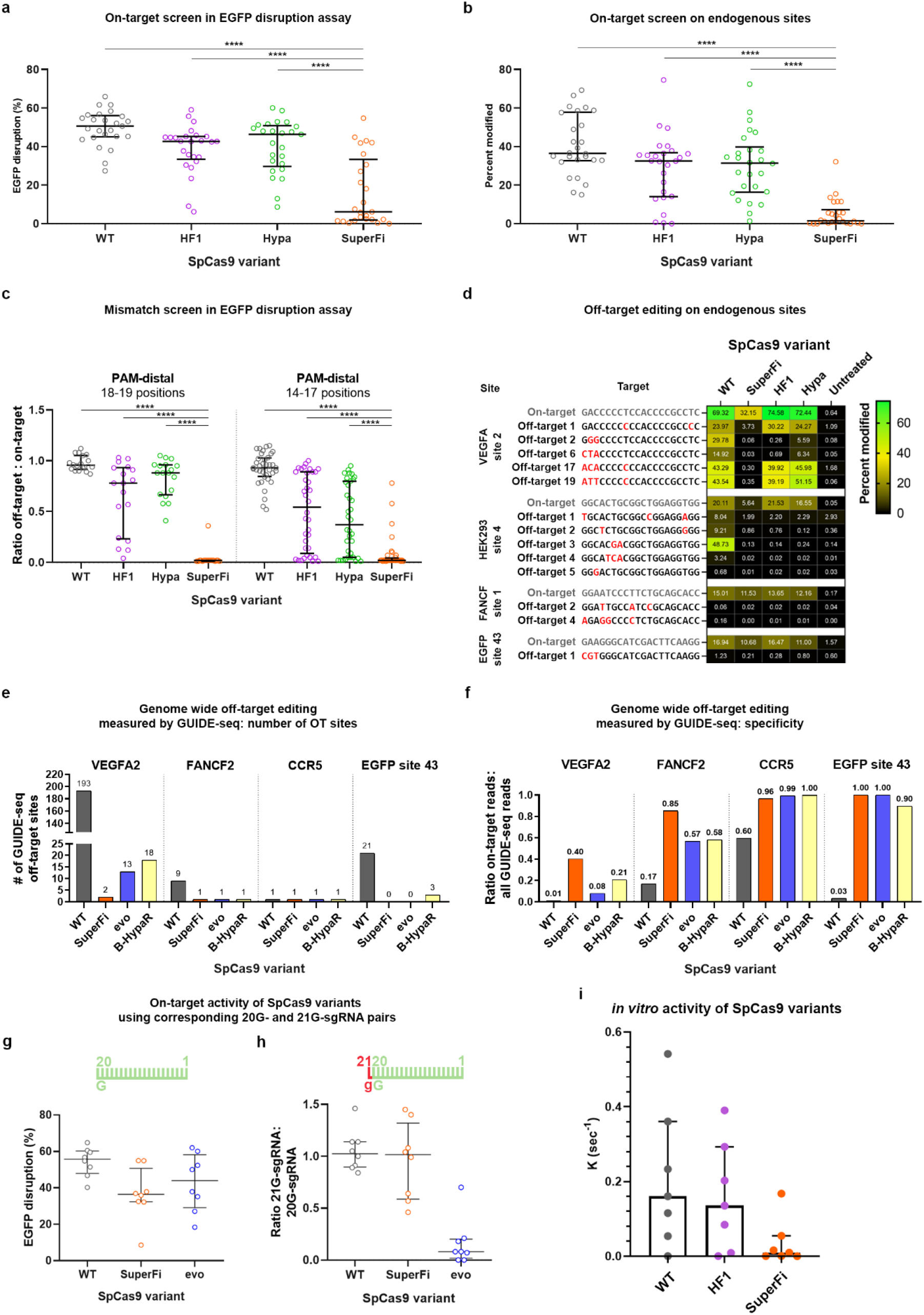
On- and off-target activities of the SuperFi-Cas9 nuclease in mammalian cells. **a-b,** On-target activity of various SpCas9 variants as measured (**a**) in EGFP disruption assay or (**b**) by NGS on endogenous target sites presented on a scatter dot plot. Data are also presented in Supplementary Figure 1b and c, respectively. **c,** Off-target activity of various SpCas9 variants as measured in EGFP disruption assay presented on a scatter dot plot. Only those data points are presented here for which the on-target activity exceeded 70%. **d,** Off-target activity of various SpCas9 variants as measured by NGS on endogenous target sites presented on a heatmap. The heatmap shows the mean on- and off-target modifications (indels) of three parallel transfections. Data related to endogenous on-target sites where editing was low (<5%) with SuperFi-Cas9 are not shown on this figure, but data is available in Supplementary Table 3. On- and off-target editing was measured from the same samples. The mismatching nucleotides at DNA off-target sites are indicated as red letters. **c-d,** Data points are from Supplementary Figure 2a and b, respectively. **e-f,** Bar charts of (**e**) the total number of off-target sites and (**f**) the on-target cleavage specificity, expressed as the percentages of the on-target reads from all reads, as measured by GUIDE-seq. Data are from Supplementary Figure 3. **g-h**, EGFP disruption activity (**g**) with 20G-sgRNAs and (**h**) the same sites targeted with 21G-sgRNAs, normalized to their 20G-sgRNA counterparts, presented on a scatter dot plot. Data are also presented in Supplementary Figure 4b. **i,** *In vitro* cleavage activities of the variants employing 7 targets of panel (a). Data is related to Supplementary Figure 8. The median and interquartile range are shown; data points represent the mean of the fitted k value triplicates. **a-c, g-h,** The median and interquartile ranges are shown; data points are plotted as open circles representing the mean of triplicates. Statistical significance was assessed by RM one-way ANOVA and statistical details and exact p-values can be found in Materials and Methods and in Supplementary Table 5.

We were curious to see if SuperFi’s strongly reduced activity could be seen in *in vitro* experiments as well. We examined its activity *in vitro*, in a DNA cleavage assay on 7 targets from Figure 1a, on which SuperFi showed WT-like (2 targets) or zero/close to zero (5 targets) activity in cells. SuperFi also showed strongly reduced activity *in vitro*, with cleavage approaching that of WT only on those targets on that also had WT-like activity in cells (Fig. 1i and Supplementary Fig. 8).

Based on the high fidelity of SuperFi, we proposed that it may be particularly effective at decreasing off-targets of base^27–30^ and prime editing^31^. We examined SuperFi in these experiments along with IFNs showing the closest activity/fidelity in a previous study^32^ concerned with the mechanism of base editors. When tested, SuperFi cytosine base editor (CBE) and SuperFi adenine base editor 7.10 (ABE7 for short) exhibited strongly reduced base editing activity, however, the activity of SuperFi-ABE8e approached that of ABE8e when assessed both in the BEAR^32^ fluorescent assay (Fig. 2a and Supplementary Fig. 5a) as well as on genomic targets (Fig. 2b and Supplementary Fig. 6a). Due to its high, undiscriminating activity, ABE8e has been used in only few applications, however, SuperFi-ABE8e successfully reduced its SpCas9-dependent off-target effects when assessed both in the BEAR assay (Fig. 2c and Supplementary Fig. 5b) as well as on genomic targets (Fig. 2d and Supplementary Fig. 6b). It also demonstrated lower level of bystander editing than ABE8e while maintaining significantly higher on-target activity than ABE7 (Fig. 2b). SuperFi in combination with a prime editor was unsuccessful, as it showed little to no activity in the PEAR^33^ fluorescent assay (Supplementary Fig. 7a) and showed no activity at all on 10 genomic loci installing 13 types of edits (Fig. 2e and Supplementary Fig. 7b).

**Figure 2.**
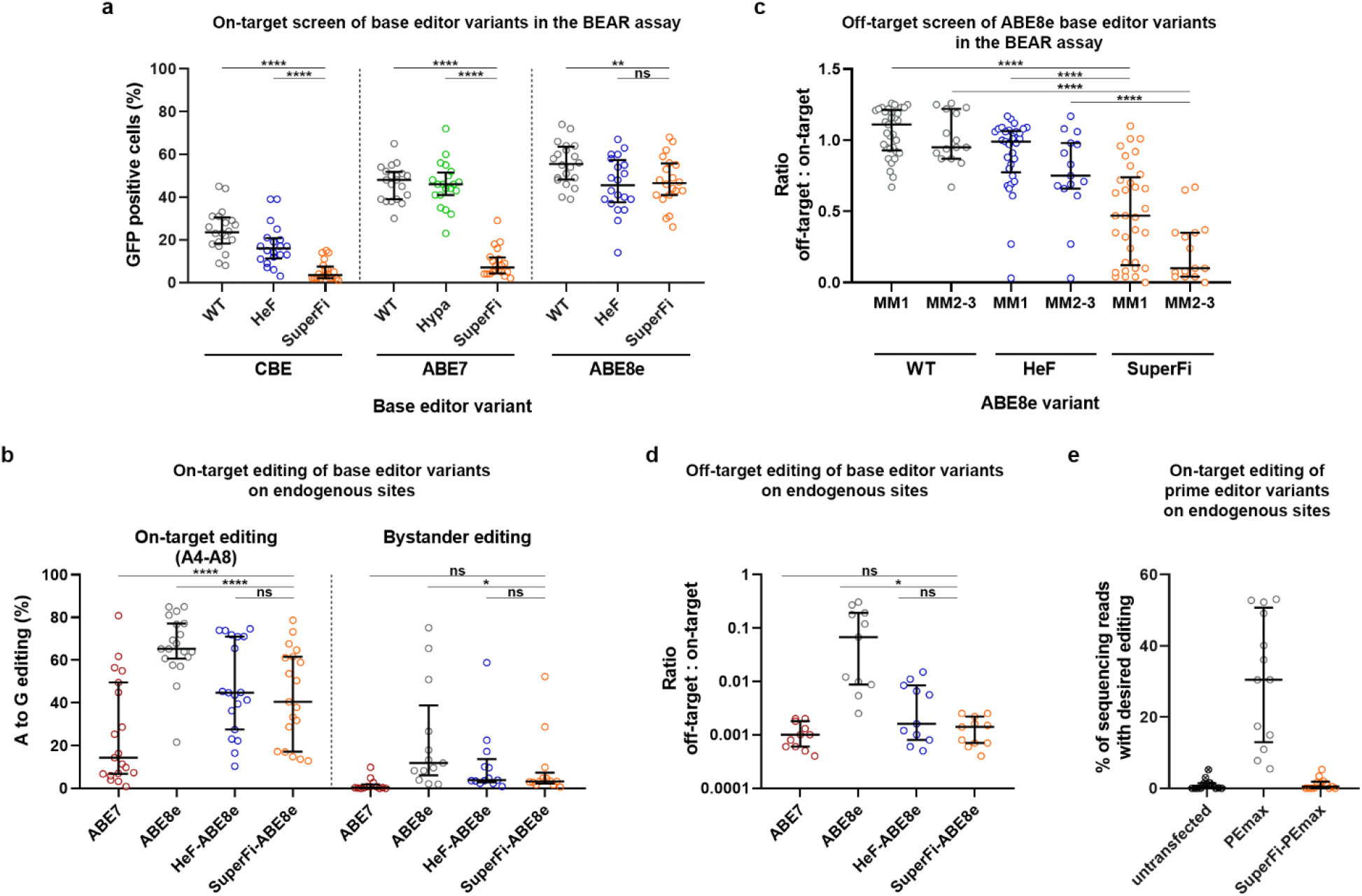
On- and off-target activities of SuperFi base and prime editors in mammalian cells. **a,** On-target base editing activity of CBE, ABE7 and ABE8e variants were assessed in the BEAR assay. Data are from Supplementary Figure 5a. **b, d,** Base editing activity of ABE8e variants on endogenous (**b**) on-target and (**d**) off-target sites as measured by NGS. Data are also presented in Supplementary Figure 6. **b**, The editing efficiency of adenines inside (A4-A8) and outside the editing window (bystander editing) is shown side by side separated with a dashed line. **d,** The relative activity (off-target/on-target ratio) of ABE variants is shown for all adenines of four genomic off-target sites. On-target adenines is the adenine with the highest editing with each on-target sequence with ABE7. **c**, The fidelity of ABE8e variants was assessed in the BEAR assay with 2 matching sgRNAs and 33 sgRNAs mismatching in one (MM1) or in two to three positions (MM2-3). The relative activity of all base editors (off-target/on-target ratio) for all adenines is plotted separately for all MM1 and all MM2-3 sgRNAs. Data are from Supplementary Figure 5b. **e,** Prime editing activity of SuperFi. Data are from Supplementary Figure 7. **a-e**, Results are presented on a scatter dot plot, the median and interquartile ranges are shown; data points are plotted as open circles representing the means of triplicates. Statistical significance was assessed by RM one-way ANOVA and statistical details and exact p-values can be found in Materials and Methods and in Supplementary Table 5.

## Discussion

Several factors could be the reason why SuperFi did not completely live up to the high expectations generated by the study of Bravo et al.^24^. One may suggest that the mutations within the RuvC loop interfere with cellular factors that specifically diminish its high *in vitro* activity in mammalian cells. The fact that SuperFi has WT-like activity on a few targets does not support this scenario. Alternatively, the single target used in ref. 24 may happen to be one of the few targets on which SuperFi can demonstrate an activity rate close to that of WT-SpCas9. The on-target activity and mismatch tolerance pattern of SuperFi is very similar to that of the highest fidelity first-generation IFNs, such as evo-, B-HypaR- and HeFSpCas9, suggesting that these mutations affect SpCas9 activity in the same way as the mutations of the first-generation IFNs^12,21,32^. Interestingly, the Blackjack mutations that remove a part of the RuvC loop including the capping 1013Tyr, also make IFNs compatible with 21G-sgRNAs similarly to SuperFi^12^. When applied without other mutations to the WT nuclease, Blackjack mutations do not decrease much of the on-target activity of SpCas9 suggesting that mutations other than that of the 1013Tyr are responsible for the large decrease in the on-target activity of SuperFi. Although its relatively low on-target activity does not make SuperFi useful for general use with various applications, in the case of the super-active ABE8e it seems to effectively counteract the over-activity of the mutant deaminase partner and becomes a very effective tool that is significantly more active than ABE7 and more specific than ABE8e.

## Materials and methods

### Materials

Restriction enzymes, T4 ligase, Dulbecco’s modified Eagle Medium DMEM (Gibco), fetal bovine serum (Gibco), Turbofect, Qubit dsDNA HS Assay Kit, Taq DNA polymerase (recombinant), Platinum Taq DNA polymerase, 0.45 μm sterile filters and penicillin/streptomycin were purchased from Thermo Fischer Scientific. DNA oligonucleotides, trimethoprim (TMP) and GenElute HP Plasmid Miniprep kit were acquired from Sigma-Aldrich. ZymoPure Plasmid Midiprep and Maxiprep kits were purchased from Zymo Research. NEBuilder HiFi DNA Assembly Master Mix and Q5 High-Fidelity DNA Polymerase were obtained from New England Biolabs Inc. NucleoSpin Gel and PCR Clean-up kit was purchased from Macherey-Nagel. SF Cell Line 4D-Nucleofector X Kit S were purchased from Lonza, Bioruptor 0.5 ml Microtubes for DNA Shearing from Diagenode. Agencourt AMPure XP beads were purchased from Beckman Coulter. T4 DNA ligase (for GUIDE-seq) and end-repair mix were acquired from Enzymatics. KAPA universal qPCR Master Mix was purchased from KAPA Biosystems.

### Plasmid construction

SuperFi-Cas9 vectors were constructed using NEBuilder HiFi DNA Assembly. All SpCas9, base and prime editor variants were subcloned to their corresponding expression plasmid backbones. For detailed cloning and sequence information see Supplementary Information. sgRNA coding plasmids were constructed as detailed in Supplementary Information. The list of sgRNA target sites, mismatching sgRNA sequences and plasmid constructs used in this study are available in Supplementary Table 1. The sequences of all plasmid constructs were confirmed by Sanger sequencing (Microsynth AG).

In all experiments the following plasmids were used for SpCas9, base and prime editor variant expression (Addgene# provided): pX330-Flag-WT_SpCas9 (without sgRNA; with silent mutations) (#126753), pX330-Flag-SpCas9-HF1 (without sgRNA; with silent mutations) (#126755), pX330-Flag-HypaSpCas9 (without sgRNA; with silent mutations) (#126756), pX330-Flag-evoSpCas9 (without sgRNA; with silent mutations) (#126758)^12^, B-HypaR-SpCas9 (#126764) (personal communication), pAT9676-ABE (#162997) for ABE7, pAT9993-Hypa-ABE (#163001) for Hypa-ABE7; pAT9675-CBE (#163007) for CBE; pAT15069-HeF-CBE (#163007) for HeF-CBE; pAT15482_ABE8e (#174120) for ABE8e; pAT15488_HeF-ABE8e (#174126) for HeF-ABE8e^32^ and pCMV-PEmax^34^ (#174820) for PEmax expression. pET-Cas9-NLS-6xHis (Addgene #62933)^7^, pET-SpCas9-HF1-NLS-6xHis, pET-SuperFi-Cas9-NLS-6xHis were used for WT, -HF1 and SuperFi-SpCas9 bacterial expression, respectively.

Plasmids developed by us in this study and deposited at Addgene are the following: pPIK16045_pX330_Flag-SuperFi-Cas9 (#184370), pAT15542_nCBE-SuperFi-Cas9 (#184372), pAT15544_nABE-SuperFi-Cas9 (#184374), pAT15546_nABE8e-SuperFi-Cas9 (#184376), pAT 15547_PEmax-SuperFi-Cas9 (#184377).

### Cell culturing and transfection

Cells employed in this study are HEK293 (Gibco 293-H cells), N2a.dd-EGFP (a neuro-2a mouse neuroblastoma cell line developed by us containing a single integrated copy of an EGFP-DHFR[DD] [EGFP-folA dihydrofolate reductase destabilization domain] fusion protein coding cassette originating from a donor plasmid with 1,000 bp long homology arms to the *Prnp* gene driven by the *Prnp* promoter (*Prnp*. HA-EGFP-DHFR[DD]), N2a.EGFP and HEK-293.EGFP (both cell lines containing a single integrated copy of an EGFP cassette driven by the *Prnp* promoter)^21^ cells. Cells were grown at 37 °C in a humidified atmosphere of 5% CO_2_ in high glucose Dulbecco’s Modified Eagle medium (DMEM) supplemented with 10% heat inactivated fetal bovine serum, 4 mM L-glutamine (Gibco), 100 units/ml penicillin and 100 μg/ml streptomycin. Cells were passaged up to 20 times (washed with PBS, detached from the plate with 0.05% Trypsin-EDTA and replated). After 20 passages, cells were discarded. Cell lines were not authenticated as they were obtained directly from a certified repository or cloned from those cell lines. Cells were tested for mycoplasma contamination.

Cells were plated in case of each cell line one day prior to transfection in 48-well plates at a density of 3 x 10^4^ cells/well. The following amounts of plasmids were mixed with 1□μL Turbofect reagent in 50□μL serum-free DMEM and were incubated for 25-30 minutes prior to being added to the cells: (1) in case of SpCas9 cleavage (either in EGFP disruption or in genomic editing): SpCas9 variant expression plasmid (137 ng) (in case of titration the amount was always supplemented with dead NmCas9 expression plasmid to 137 ng) and sgRNA and mCherry coding plasmid (97 ng); (2) in case of genomic base editing: base editor variant expression plasmid (190ng) and sgRNA and BFP coding plasmid (83 ng); (3) in case of genomic prime editing: prime editor variant expression plasmid (222ng), pegRNA and BFP coding plasmid (55 ng), and second nicking sgRNA and BFP coding plasmid (36ng).

EGFP disruption experiments were conducted in N2a.EGFP cells for the on-target screen. In this cell line the EGFP disruption level is not saturated, this way this assay is a more sensitive reporter of the intrinsic activities of these nucleases compared to N2a.dd-EGFP cell line. EGFP disruption experiments were conducted in N2a.EGFP cells for the mismatch screen, 21G- and truncated sgRNA screen. In this cell line four days post-transfection results show a close to saturated level, this way it is a good reporter system for seeing the full spectrum of activities.^12,21^

The BEAR and PEAR reporters were used as described previously^32,33^. Briefly, cells were transfected as described above where in case of BEAR experiments: 66□ng of BEAR target plasmid, 56 ng of sgRNA-mCherry and 127□ng of base editor coding plasmid were used. In case of PEAR experiments: 40□ng of PEAR-GFP-2in1 target plasmid, 56 ng of mock pegRNA-BFP (to follow transfection efficiency) and 255 □ng of prime editor coding plasmid were used.

Cells were analysed by flow cytometry three days post-transfection in case of BEAR and PEAR experiments and four days post-transfection in all other experiments. Transfections were performed in triplicates. Transfection efficacy was calculated via mCherry or BFP expression. Data of the EGFP disruption, BEAR and PEAR experiments are available in Supplementary Table 2.

### Electroporation in GUIDE-seq experiments

Briefly, 2 x 10^5^ cells were resuspended in transfection solution (see below) and mixed with 666 ng of SpCas9 variant expression plasmid and 334 ng of sgRNA and mCherry coding plasmid and an additional 30 pmol dsODN (according to the original GUIDE-seq protocol^25^) was added to the mixture. Nucleofections were performed in the case of HEK293 and HEK-293.EGFP cell lines using the CM-130 program on a Lonza 4-D Nucleofector instrument on strip with 20 μl SF solution according to the manufacturer’s protocol. Transfections were performed in triplicates. Transfection efficacy was calculated via mCherry expression.

### Flow cytometry

Flow cytometry analyses were carried out on an Attune NxT Acoustic Focusing Cytometer (Applied Biosystems). For data analysis Attune NxT Software v.4.2 was used. Viable single cells were gated based on side and forward light-scatter parameters and a total of 5,000 to 10,000 viable single cell events were acquired in all experiments. BFP, GFP and mCherry signals were detected using the 405 (for BFP), 488 (for GFP) and 561 nm (for mCherry) diode laser for excitation, and the 440/50 (BFP), 530/30 (GFP) and 620/15 (mCherry) filter for emission.

### Indel, base and prime editing analysis by next-generation sequencing (NGS)

Transfected cells were analysed by flow cytometry followed by genomic DNA purification according to the Puregene DNA Purification protocol (Gentra systems). Amplicons for next generation sequencing were generated using two rounds of PCR to attach Illumina handles from the genomic DNA samples. The 1^st^ step PCR primers used to amplify target genomic sequences are listed in Supplementary Table 1: NGS primers. PCR was done in a S1000 Thermal Cycler (Bio-Rad) or PCRmax Alpha AC2 Thermal Cycler using Q5 high-fidelity polymerase supplemented with Q5 buffer (in case of *VEGFA* site 2 amplicon supplemented with Q5 High GC enhancer as well) and 150 ng of genomic DNA in a total volume of 25 μl. The thermal cycling profile of the PCR was: 98 °C 30 sec; 35 x (denaturation: 98 °C 20 sec; annealing: see Supplementary Table 1: NGS primer, 30 sec; elongation: 72 °C, see Supplementary Table 1: NGS primer); 72°C 5 min. i5 and i7 Illumina adapters were added in a second PCR reaction using Q5 high-fidelity polymerase with supplied Q5 buffer (in case of *VEGFA* site 2 amplicon together with Q5 High GC enhancer) and 1 μl of first step PCR product in total volume of 25 μl. The thermal cycling profile of the PCR was: 98 °C 30 sec; 35 x (98 °C 20 sec, 67 °C 30 sec, 72 °C 20 sec); 72°C 5 min. Amplicons were purified by agarose gel electrophoresis. Samples were quantified with Qubit dsDNA HS Assay kit and pooled. Double-indexed libraries were sequenced on a MiniSeq or NextSeq (Illumina) giving paired-end sequences of 2 x 150 bp, performed by Deltabio Ltd. Reads were aligned to the reference sequence using BBMap.

Indels were counted computationally amongst the aligned reads that matched at least 75% of the first 20□bp of the reference amplicon. Indels without mismatches were searched starting at ±2□bp around the cut site. For each sample, the indel frequency was determined as (number of reads with an indel) / (number of total reads). Frequency of substitutions without indels generated by base or prime editing was determined as the percentage of (sequencing reads with the intended modification, without indels) / (number of total reads). By contrast, frequency of intended insertions or deletions generated by prime editing was determined as the percentage of (all sequencing reads with the intended modification) / (number of total reads). For these samples the indel background was calculated from reads containing types of indels that were different from the aimed edit.

The following software were used: BBMap 38.08, samtools 1.8, BioPython 1.71, PySam 0.13. For NGS data information see Supplementary Table 3. The deep sequencing data have been submitted to the NCBI Sequence Read Archive and will be available upon publication.

### *In vitro* transcription

sgRNAs were transcribed *in vitro* using TranscriptAid T7 High Yield Transcription Kit and PCR-generated double-stranded DNA templates carrying a T7 promoter sequence. PCR primers used for the preparation of the DNA templates are listed in Supplementary Table 1. sgRNAs were purified with the RNA Clean & Concentrator kit and reannealed (95 °C for 5 min, ramp to 25 °C at 0.3 °C/s). sgRNAs were quality checked using 10% denaturing polyacrylamide gels and ethidium bromide staining.

### Protein purification

WT SpCas9 was purified using pET-Cas9-NLS-6xHis (Addgene #62933)^7^ plasmid, SpCas9-HF1 and SuperFi-SpCas9 were subcloned into that plasmid (details in Methods: Plasmid construction section and in Supplementary Information). The expression constructs of the SpCas9 variants were transformed into *E. coli* BL21 Rosetta 2 (DE3) cells, grown in Luria-Bertani (LB) medium at 37 °C for 16 h. 10 ml from this culture was inoculated into 1 l of growth media (12 g/l Tripton, 24 g/l Yeast, 10 g/l NaCl, 883 mg/l NaH_2_PO_4_ H_2_O, 4.77 g/l Na_2_HPO_4_, pH 7.5) and cells were grown at 37 °C to a final cell density of 0.6 OD600, and then were cooled to 18 °C. The protein was expressed at 18 °C for 16 h following induction with 0.2 mM IPTG. Proteins were purified by a combination of chromatographic steps by NGC Scout Medium-Pressure Chromatography Systems (Bio-Rad). The bacterial cells were centrifuged at 6,000 rcf for 15 min at 4 °C. The cells were resuspended in 30 ml of Lysis Buffer (40 mM Tris pH 7.5, 500 mM NaCl, 20 mM imidazole, 0.5 mM TCEP) supplemented with Protease Inhibitor Cocktail (1 tablet/30 ml; complete, EDTA-free, Roche) and sonicated on ice. Lysate was cleared by centrifugation at 48,000 rcf for 40 min at 4 °C. Clarified lysate was bound to a 5 ml HisTrap™ High Performance Ni-Charged column (Cytiva). The resin was washed extensively with a solution of 40 mM Tris pH 7.5, 500 mM NaCl, 20 mM imidazole, and the bound proteins were eluted by a solution of 40 mM Tris pH 7.5, 250 mM imidazole, 150 mM NaCl.. 10% glycerol was added to the eluted sample. The volume of the protein solution was made up to 60 ml with buffer (20 mM HEPES pH 7.5, 100 mM KCl, 1 mM DTT). Proteins were purified on a 5 ml HiTrap SP HP cation exchange column (GE Healthcare) and eluted with 1 M KCl, 20 mM HEPES pH 7.5, 1 mM DTT. They were then further purified by size exclusion chromatography on a Superdex 200 10/300 GL column (GE Healthcare) in 20 mM HEPES pH 7.5, 200 mM KCl, 1 mM DTT and 10% glycerol. The eluted protein was confirmed by SDS-PAGE and Coomassie brilliant blue R-250 staining, and they were stored at −20 °C.

### Determining active SpCas9 quantity in solution

The quantification method was based on Liu et al.^74^. The quantity of active SpCas9 protein in solution was determined using EGFP target site 5, that has shown high cleavage activity with all three proteins tested based on previous experiments. The measurement procedure is as follows: The target plasmid was incubated for an hour with protein-sgRNA complex, in different concentrations. Concentrations were determined by spectrophotometry (Nanodrop OneC) and then the target site containing plasmid (6.2 nM) and the SpCas9 protein were mixed with a ratio between 1:0.8 and 1:12 while the quantity of the sgRNA was twice that of the protein in each case. To terminate cleavage reaction, the EDTA solution (final concentration: 50 mM) was added to the reaction mix at 70 °C. Samples were ran on a 1% agarose gel. Following densitometry (GelQuantNET, BiochemLabSolutions.com), the ratio of intact plasmid and total DNA was calculated for each sample. These values were plotted and fitted on a ‘One-phase exponential decay function with time constant parameter’ curve in Origin 2018. Taken the results of this experiment, the active SpCas9 variant quantities in solution were calculated. It was also taken into consideration that SpCas9 has a one-fold turnover rate.

### Determining cleavage rate of SpCas9 variants *in vitro*

At first, two different solutions were made: (1) target plasmid solution and (2) an SpCas9-sgRNA master mix. After mixing them (see below) the ratio of the target site containing plasmid and active protein was 1:3. Both solutions were diluted with the same cleavage buffer (final concentration: 20 mM HEPES pH 7.5, 200 mM KCl, 1 mM TCEP, 2% glycerol) and were pre-incubated at 37 °C before rmixing. To trigger cleavage reaction, the target plasmid solution was added to the SpCas9-sgRNA mixture. To terminate cleavage reaction EDTA (final concentration: 50 mM) was added to the reaction mix at 70 °C in different time points. First, we determined the approximate cleavage rate for every protein-target combination. Based on these preliminary results, we have defined three categories for the time range in which sampling should take place: fast (3-30 sec), medium (3-300 sec), and slow (3-3600 sec). To record precisely the actual time of the sampling a digital chronometer was attached to the pipette which can record time points in an application developed by us. Samples were then run on a 1% agarose gel. Following densitometry (GelQuantNET, BiochemLabSolutions.com), the cleaved DNA was calculated for each sample. Experiments were performed in triplicates. These values were plotted and fitted on a ‘One-phase exponential decay function with time constant parameter’ curve in Origin 2018. All fitted curves are available in Supplementary Figure 8, the k values are available in Supplementary Table 6.

### GUIDE-seq

In the first step genomic DNA was sheared with BioraptorPlus (Diagenode) to 550 bp in average. Sample libraries were assembled as previously described^25^ and sequenced on Illumina MiniSeq instrument by Deltabio Ltd. Data were analysed using open-source GUIDE-seq software (version 1.1)^35^. Consolidated reads were mapped to the human reference genome GrCh37 in case of EGFP target site 43 supplemented with the integrated EGFP sequence). Upon identification of the genomic regions integrating doublestranded oligodeoxynucleotide (dsODNs) in aligned data, off-target sites were retained if at most seven mismatches against the target were present and if absent in the background controls. Visualization of aligned off-target sites are provided as a color-coded sequence grid. Summarized results can be found in Supplementary Table 4 and GUIDE-seq sequencing data are deposited at NCBI Sequence Read Archive and will be available upon publication.

### Western blot

N2a.dd-EGFP cells were cultured on 48-well plates and were transfected as described above in the EGFP disruption assay section. Four days post-transfection, 9 parallel samples corresponding to each SpCas9 variant transfected were washed with PBS, then trypsinized and mixed, and were analysed for transfection efficiency via mCherry fluorescence level by using flow cytometry. The cells from the mixtures were centrifuged at 200 rcf for 5 min at 4 °C. Pellets were resuspended in ice cold Harlow buffer (50 mM Hepes pH 7.5; 0.2 mM EDTA; 10 mM NaF; 0.5% NP40; 250 mM NaCl; Protease Inhibitor Cocktail 1:100; Calpain inhibitor 1:100; 1 mM DTT) and lysed for 20-30 min on ice. The cell lysates were centrifuged at 19,000 rcf for 10 min. The supernatants were transferred into new tubes and total protein concentrations were measured by the Bradford protein assay. Before SDS gel loading, samples were boiled in Protein Loading Dye for 10 min at 95 °C. Proteins were separated by SDS-PAGE using 7.5% polyacrylamide gels and were transferred to a PVDF membrane, using a wet blotting system (Bio-Rad). Membranes were blocked by 5% non-fat milk in Tris buffered saline with Tween20 (TBST) (blocking buffer) for 2 h. Blots were incubated with primary antibodies [anti-FLAG (F1804, Sigma) at 1:1,000 dilution; anti-β-actin (A1978, Sigma) at 1:4,000 dilution in blocking buffer] overnight at 4 °C. The next day, after washing steps in TBST, the membranes were incubated for 1 h with HRP-conjugated secondary anti-mouse antibody 1:20,000 (715-035-151, Jackson ImmunoResearch) in blocking buffer. The signal from detected proteins was visualized by ECL (Pierce ECL Western Blotting Substrate, Thermo Scientific) using a CCD camera (Bio-Rad ChemiDoc MP).

### Statistics

Differences between SpCas9 variants were tested by using RM one-way ANOVA and Dunnett’s multiple comparisons test with a single pooled variance (Fig. 1b, 2a, 2b-on-target editing, 2c-MM1) or by using RM one-way ANOVA, with the Geisser-Greenhouse correction and Dunnett’s multiple comparisons test with individual variances computed for each comparison (Fig. 1a, 1c, 2b-bystander editing, 2c-MM2-3, 2d) in the cases where sphericity did not meet the assumptions of RM one-way ANOVA. Statistical tests were performed using GraphPad Prism 8. Test results are shown in Supplementary Table 5.

## Supplementary Figures and legends

**Supplementary Figure 1.**
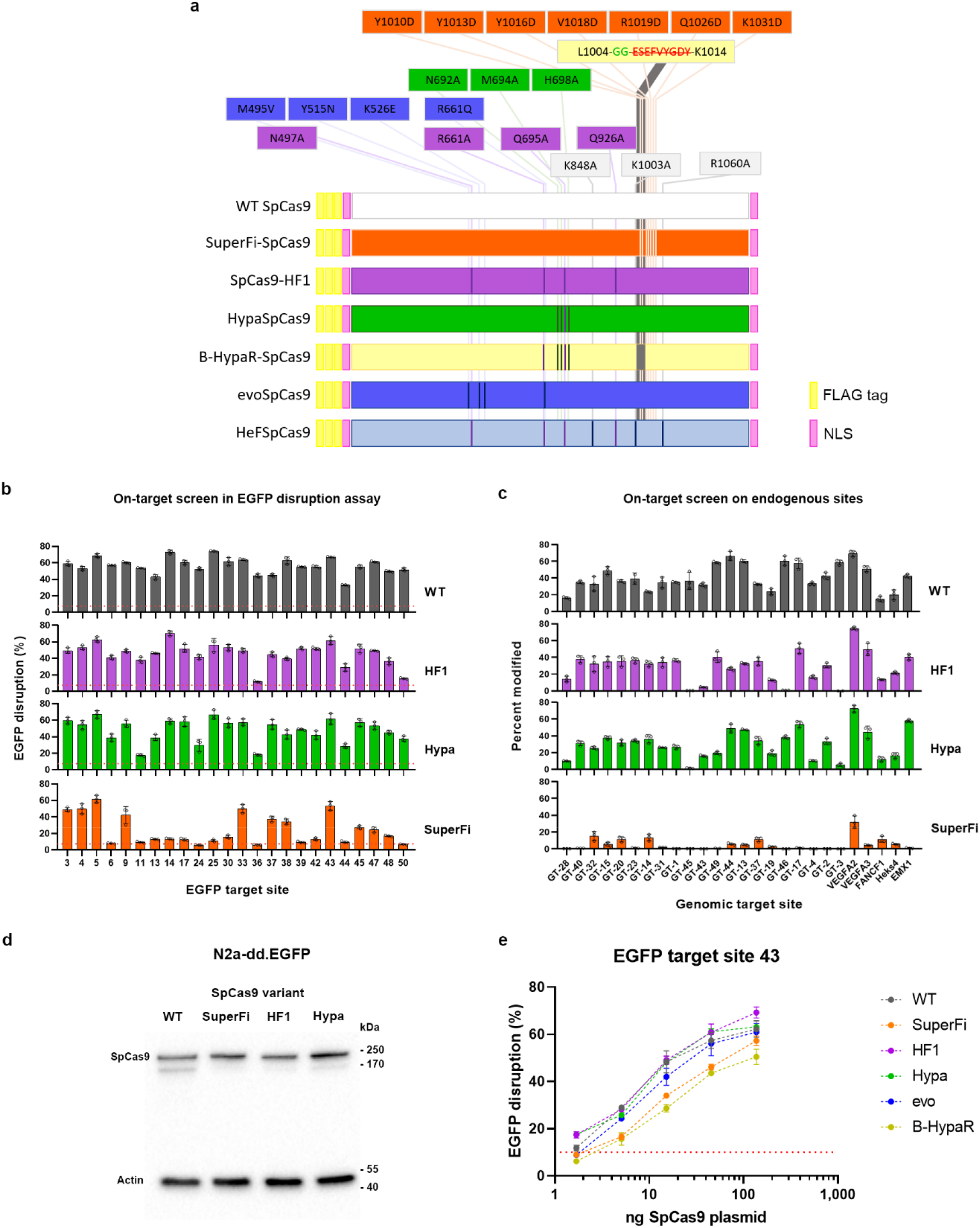
On-target activity of SuperFi-Cas9. **a,** Schematic representation of the polypeptide chain of SpCas9s. Mutations of the IFNs used in this study are indicated on the linear sequence of SpCas9. **b,** On-target EGFP disruption data of WT SpCas9 and three IFN variants, as indicated in the panels, on 24 EGFP target sites. **c,** On-target activity of WT SpCas9 and three IFN variants, as indicated in the panels, across 26 endogenous sites as measured by NGS. **b-c,** Means are shown, error bars represent the standard deviation (SD) for triplicates (overlaid as white circles). (**b**) Level of background EGFP loss is indicated by a red dashed line (average of the percentage of dead SpCas9 controls from all target sites). (**c**) In case of endogenous target sites the measured background level can be found in Supplementary Table 3. **d,** Immunoblot analysis of the expression levels of SpCas9 nuclease variants in cell lysates of reporter N2a.dd-EGFP cells transfected with the indicated nuclease constructs. β-actin was used as a loading control for total protein amounts analyzed. **e,** Titration of expression plasmid amounts of wild-type SpCas9 and different IFN variants. Means are shown in case of each data point, error bars represent the standard deviation (SD) for triplicates; mean level of background EGFP loss in negative controls is represented by the red dashed line. **b-c**, Target sequences, EGFP disruption, NGS data and statistical details are reported in Supplementary Table 1-3 and 5.

**Supplementary Figure 2.**
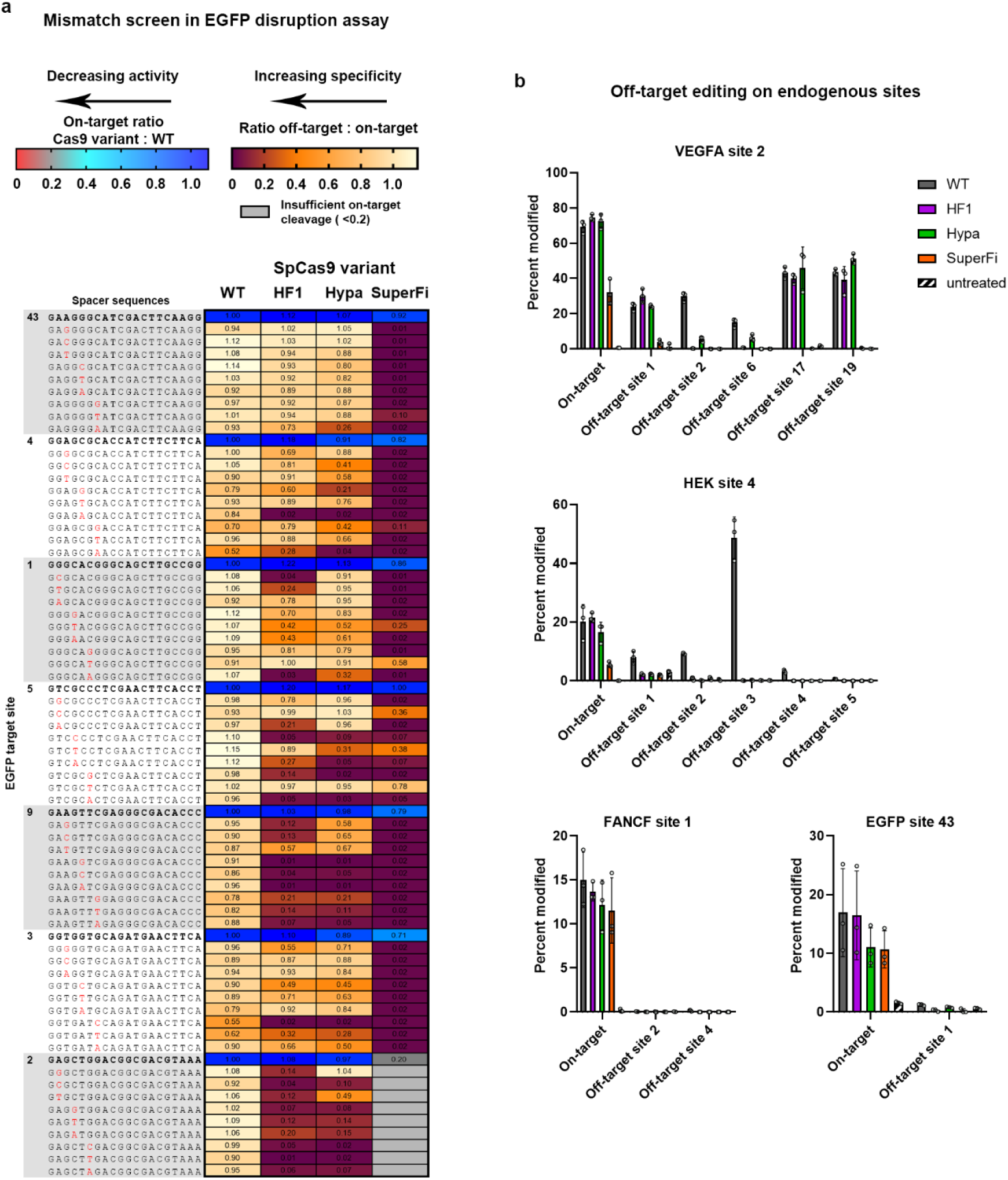
Mismatch tolerance of SuperFi-Cas9. **a,** Mismatch screen of nuclease variants with either perfectly matching sgRNAs (red to blue) or single mismatching sgRNAs (yellow to brown) presented on a heatmap. Grey boxes: not determined due to too low on-target activity. **b,** On-target activity of WT SpCas9 and three IFN variants, as indicated in the panels, across 4 endogenous on-target sites with a total of 13 off-target sites as measured by NGS. Off-target sites were selected from previous GUIDE-seq experiments^12,25^. Means are shown, error bars represent the standard deviation (SD) for triplicates (overlaid as white circles). **a-b**, Target sequences, EGFP disruption, NGS data and statistical details are reported in Supplementary Table 1-3 and 5.

**Supplementary Figure 3.**
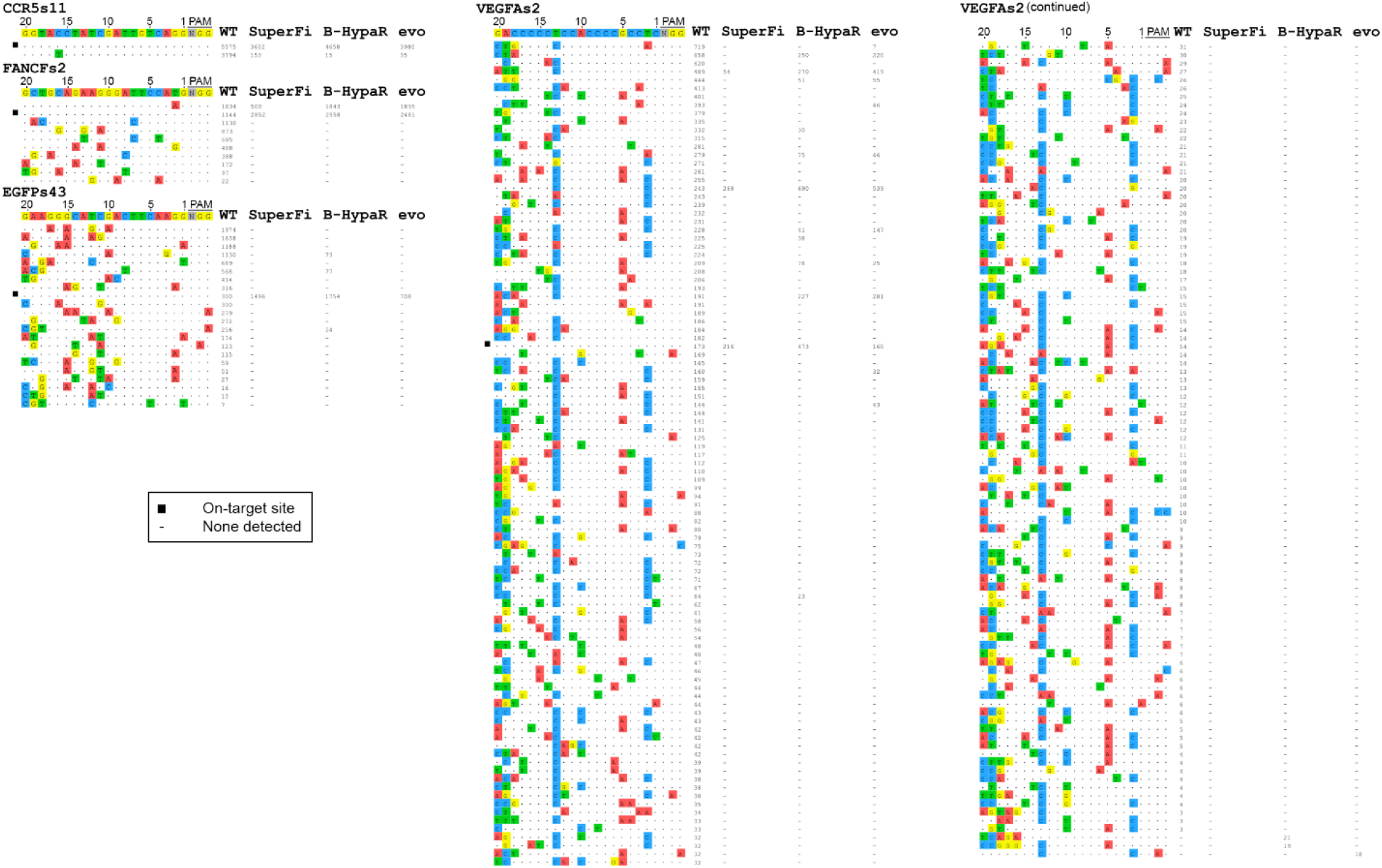
Genome-wide off-target effects of SuperFi-Cas9 variant identified by GUIDE-seq. Off-target cleavage sites of SpCas9 variants identified by GUIDE-seq. Read counts represent a measure of cleavage frequency at a given site; mismatched positions within the spacer or PAM are highlighted in different colours. (-) indicates zero reads, which means that off-target cleavage was not detected, black squares indicate the on-target sites. Target sequences and GUIDE-seq data are available in Supplementary Table 1 and 4.

**Supplementary Figure 4.**
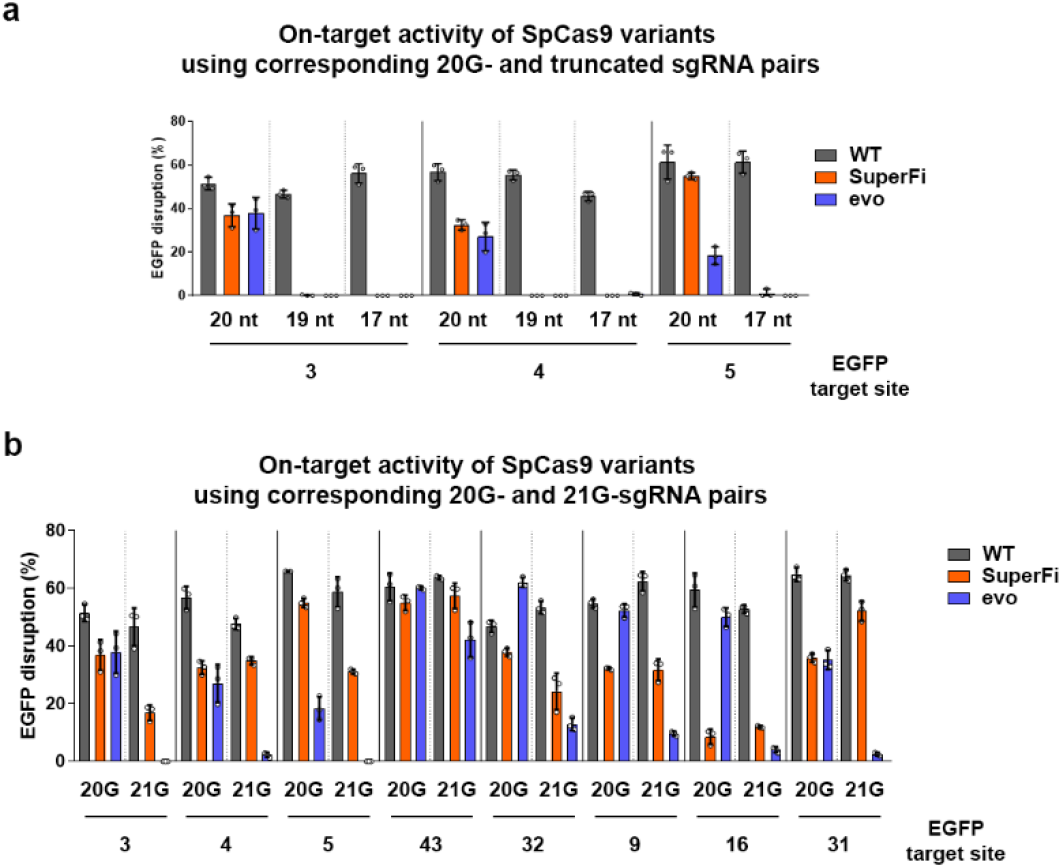
Tolerance of SuperFi-Cas9 to 5’ modified sgRNAs. Effects of (**a**) 5’ truncation of sgRNAs and (**b**) 21G-sgRNAs measured on three and eight targets, respectively, on the activity of WT and three SpCas9 variants as indicated in the panels. Means are shown, error bars represent the standard deviation (SD) for triplicates (overlaid as white circles). Target sequences, EGFP disruption and statistical details are reported in Supplementary Table 1, 2 and 5.

**Supplementary Figure 5.**
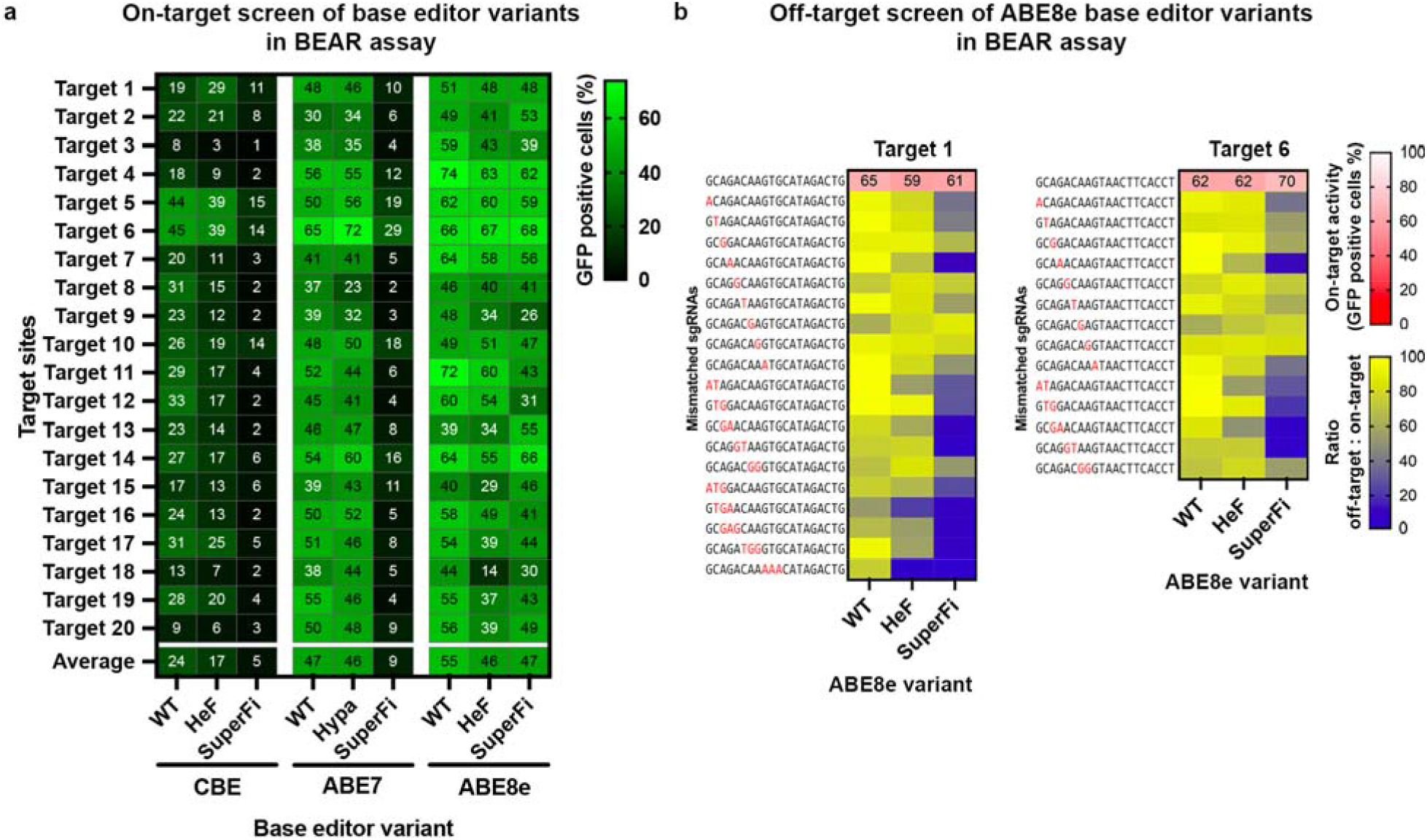
On- and off-target activities of SuperFi base editor variants in the BEAR assay. **a**, On-target activity of WT and base editor variants as indicated in the panel. The heatmap shows mean on-target base editing activity (as GFP positive cells) of triplicates on 20 different target sites. **b,** Mismatch tolerance of WT-ABE8e and IFN base editor variants as indicated in the panel were examined using 35 sgRNAs on two targets (Target 1 and 6 from panel [**a]**) with perfectly matching sgRNAs or sgRNAs mismatching in one, two or three positions as indicated by red letters in the spacer sequences. White and red heatmaps show the mean on-target activity (percentages of GFP positive cells) derived from triplicates. Blue and yellow heatmaps show the ratio of off-target/on-target activity derived from triplicates. Target sequences, BEAR assay data and statistical details are reported in Supplementary Table 1, 2 and 5.

**Supplementary Figure 6.**
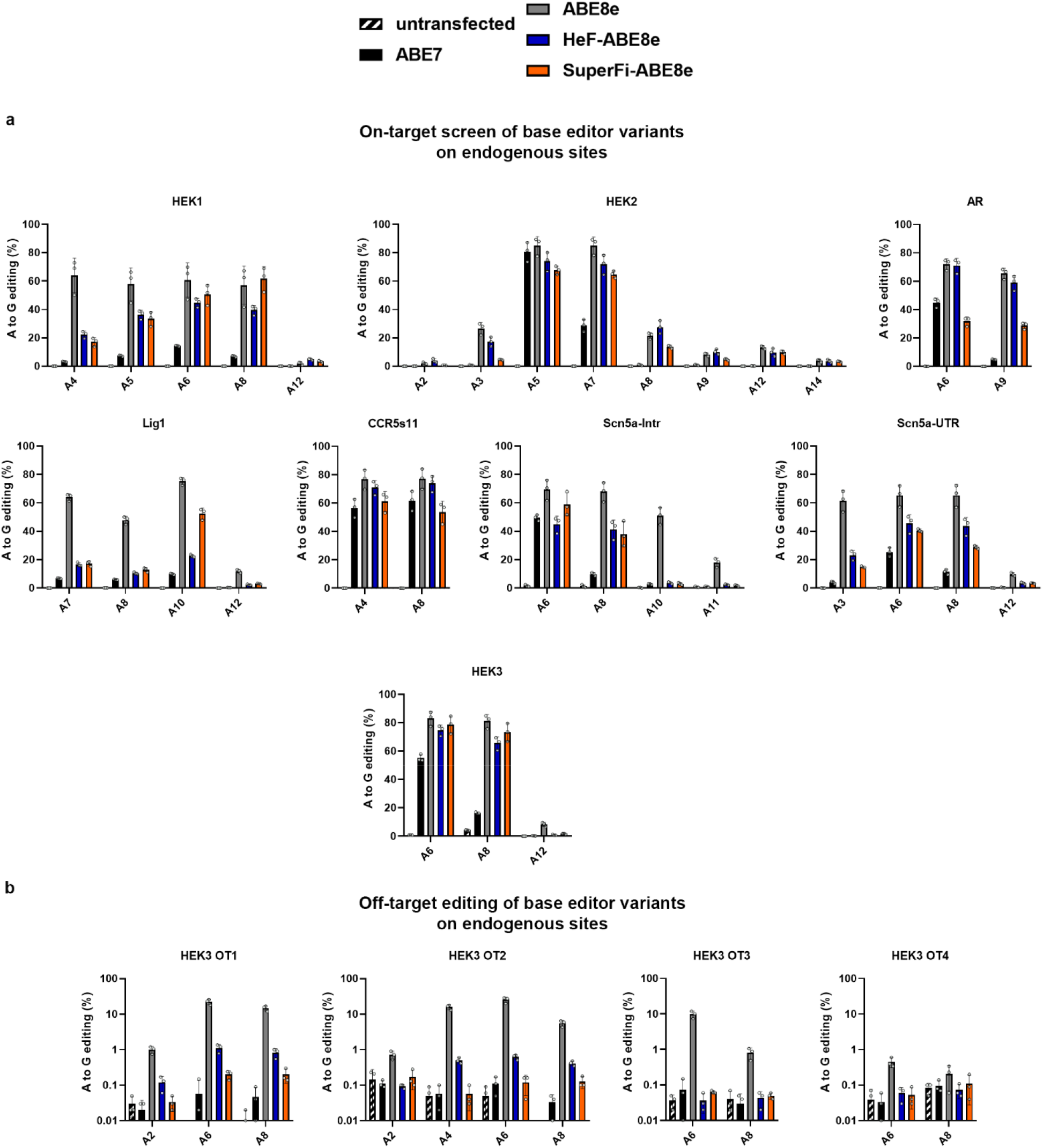
On- and off-target activities of SuperFi adenine base editor variant on endogenous target sites. Editing efficiencies of ABE7, WT-ABE8e, and two IFN ABE8e base editor variants are shown (**a**) on 8 genomic loci for assessing on-target and (**b**) on 4 loci for assessing off-target base editing as measured by NGS. **a, b,** The diagrams show A to G conversion ratios at each adenine position for each target. As negative controls, base conversion was measured for untransfected cells (black with white dashes). Means are shown, error bars represent the standard deviation (SD) for triplicates (overlaid as white circles). Target sequences, NGS data and statistical details are reported in Supplementary Table 1, 3 and 5.

**Supplementary Figure 7.**
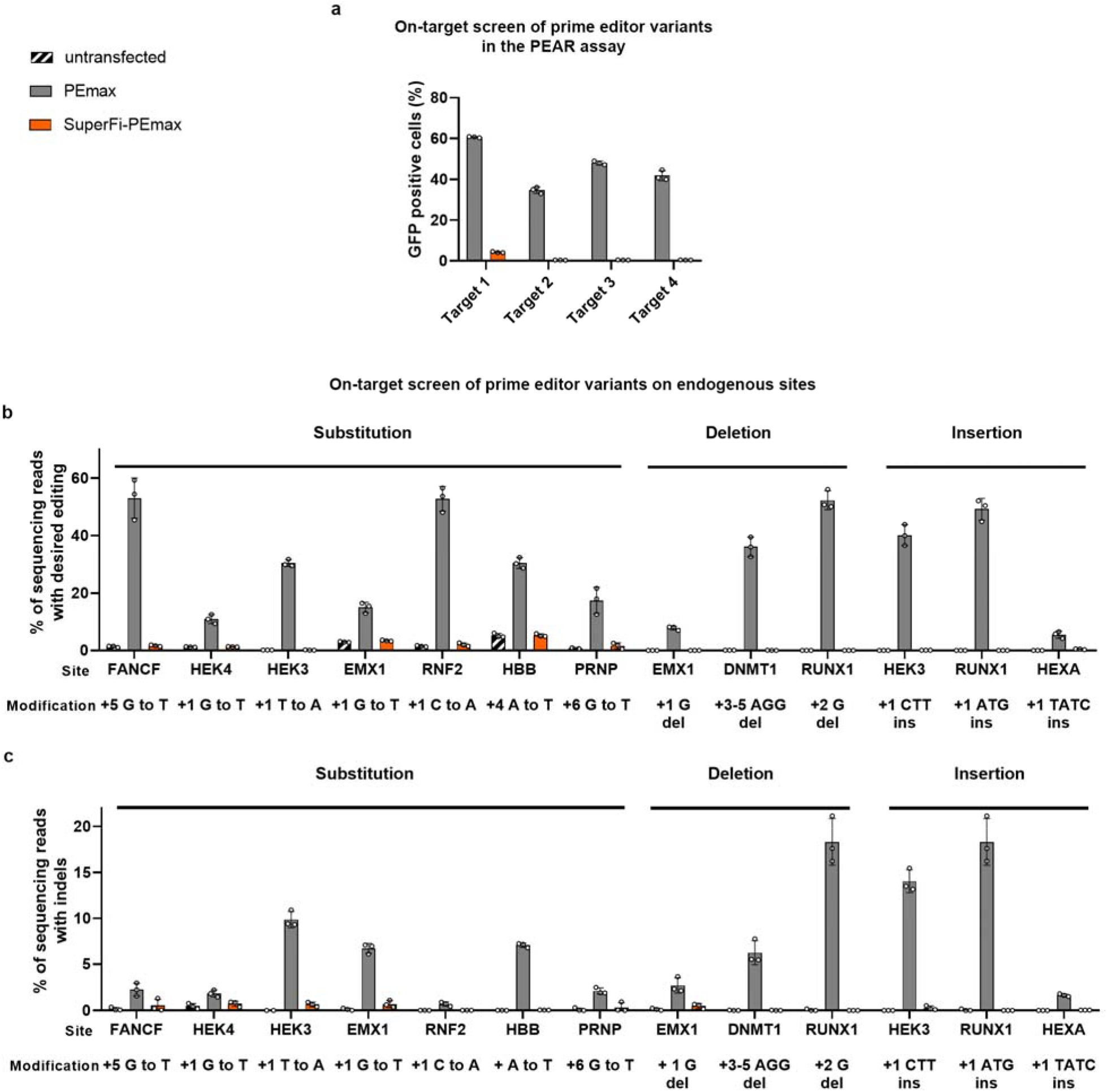
Editing activities of the SuperFi prime editor variant. **a,** Prime editing efficiencies of PEmax and SuperFi-PEmax, as indicated in the panel, were assessed in the PEAR fluorescent assay exploiting a pegRNA-target plasmid (PEAR-GFP-2in1)^33^ bearing four different spacer and target sequences. **b**, Prime editing efficiencies (desired editing without indels) and **c**, unwanted indels of PEmax and SuperFi-PEmax were measured on 10 genomic loci installing 13 edits including substitutions, insertions and deletions, as assessed by NGS. As negative controls, base conversions are also shown for untransfected cells (black with white dashes). Means are shown, error bars represent the standard deviation (SD) for triplicates (overlaid as white circles). Target sequences, PEAR assay, NGS data and statistical details are reported in Supplementary Table 1-3 and 5.

**Supplementary Figure 8:**
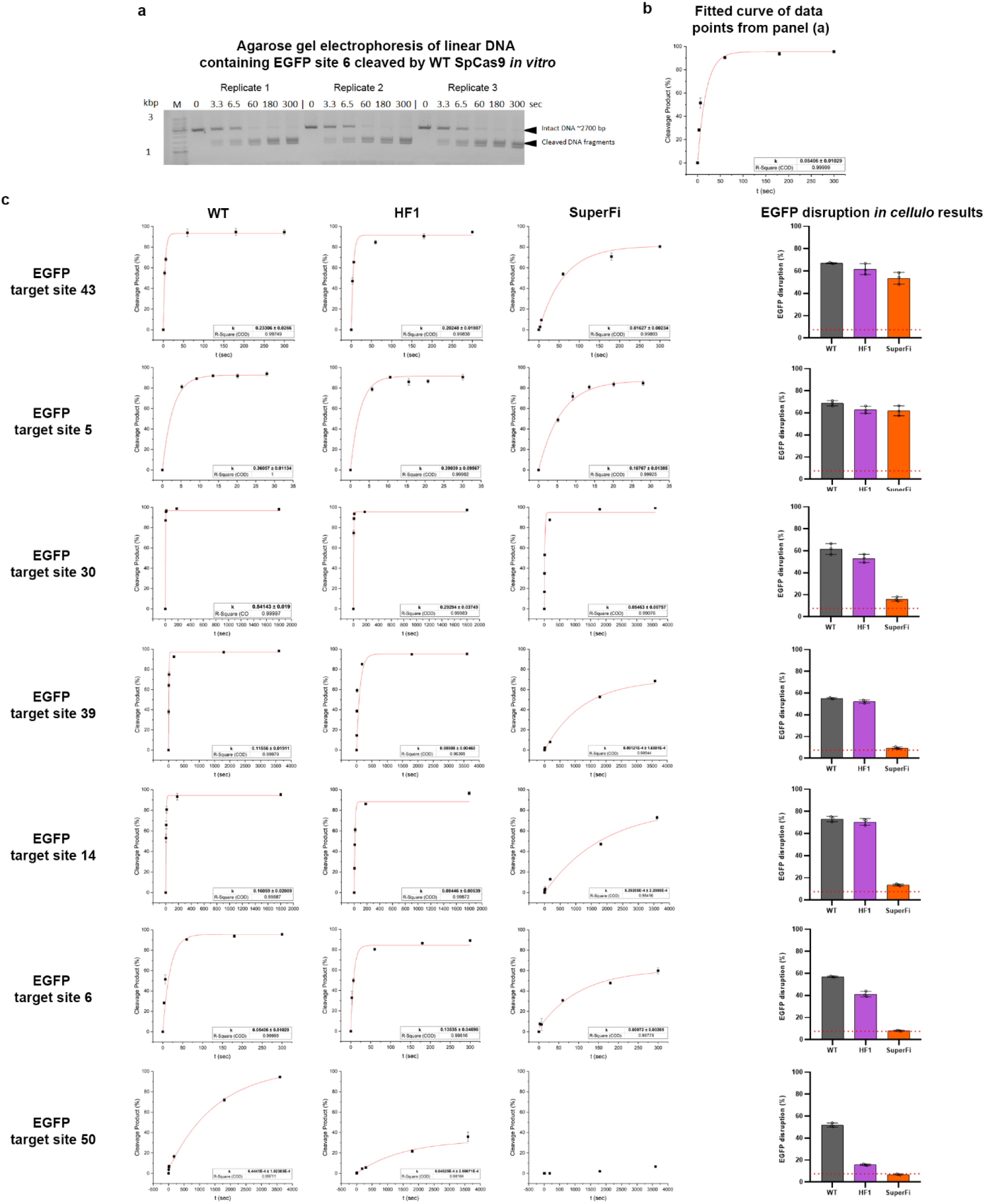
*In vitro* fitted curves on 7 EGFP target sites cleaved with WT, -HF1 or SuperFi-SpCas9 variants. **a,** Representative agarose gel showing the activity of WT SpCas9 in a linear DNA-cleaving *in vitro* assay at different timepoints in triplicates. **b,** Plot showing values representing the amount of the cleaved DNA derived from the intensity of bands from the representative agarose gel in panel (a). **c,** Plots show values derived from the band intensities measured on agarose gel for all 7 target sites. Exponential curves were fitted to the fraction of cleaved DNA measured during the time frame of the experiments in triplicates. The values and average K values were derived from these fitted curves. On the right side of the panel EGFP disruption data from Figure 1a (and Supplementary Figure 1b) is shown for each target site. Summary of target and primer sequences and *in vitro* data are reported in Supplementary Table 1 and 6.

## Data availability

Expression vectors developed in this study are available from Addgene: pPIK16045_pX330_ Flag-SuperFi-Cas9 (#184370), pAT15542_nCBE-SuperFi-Cas9(#184372), pAT15544_ nABE-SuperFi-Cas9 (#184374), pAT15546_nABE8e-SuperFi-Cas9 (#184376), pAT15547_ PEmax-SuperFi-Cas9 (#184377). The deep sequencing data have been submitted to the NCBI Sequence Read Archive and will be available upon publication.

## Acknowledgements

We thank Ildikó Szűcsné Pulinka, Judit Szűcs, Vivien Karl, Ferencné Zájer and Diána Szeregnyei for their excellent laboratory assistance. We thank Sarah Laura Krausz for NGS bioinformatics, Vanessza Laura Végi for proofreading the manuscript and for Zsuzsa Bartos and Zsófia Rakvács for their valuable help.

## Author contributions

P.I.K and A.T. contributed equally to this work. P.I.K and A.T. performed the experiments, processed the data and presented the results. All authors designed the experiments and interpreted the results. E.W. wrote the manuscript with input from the other authors.

