## Supplementary Information for "SuperFi-Cas9 exhibits extremely high fidelity but reduced activity in mammalian cells"

Summary

[*Prnp*.HA-EGFP-DHFR[DD]	16](#_Toc108439092)

### Supplementary Sequences

For detailed primer, oligonucleotide, Addgene number and SpCas9 construct information see Supplementary Table 1. The sequences of all plasmid constructs were confirmed by Sanger sequencing.

#### Cas9 spacer cloning

sgRNA expression plasmids were constructed by ligating annealed DNA oligonucleotides harboring the spacer sequence with 4 nt-long overhangs into a *Bbs*I restriction enzyme digested pmCherry_gRNA (#80457), or pmCherry_gRNA_ver2 (Addgene #126776; this plasmid backbone lacks a truncated extra guideRNA scaffold sequence) plasmids^1^. A one-pot digestion-ligation protocol was followed^2^. The synthetic DNA oligonucleotides were hybridized and the annealed oligonucleotides (2.5 µM) with 50 ng plasmid, 3 units of *Bbs*I restriction enzyme and 1.5 units of T4 DNA ligase were mixed in Green buffer (Thermo Fisher Scientific) containing 500 µM ATP. The mixture was kept at 37 °C for one hour before transforming into chemically competent Stable Competent *E. coli* cells (NEB). Two single colonies, formed after culturing on an agar plate, were tested by restriction enzyme digestion and appropriate clones were sent for sequencing.

#### SpCas9 variants, human expression plasmids

##### pX330-Flag-WT SpCas9 (Addgene #126753)

All SpCas9 variant coding plasmids’ backbone are identical with the hereunder shown sequence.

Whole DNA sequence of the plasmid:

Human codon optimized *S. pyogenes* Cas9 coloured in purple, NLS underlined, 3xFLAG tag in *light blue*, silent mutations underlined and marked with yellow*.*

ctagaggtacccgttacataacttacggtaaatggcccgcctggctgaccgcccaacgacccccgcccattgacgtcaatagtaacgccaatagggactttccattgacgtcaatgggtggagtatttacggtaaactgcccacttggcagtacatcaagtgtatcatatgccaagtacgccccctattgacgtcaatgacggtaaatggcccgcctggcattgtgcccagtacatgaccttatgggactttcctacttggcagtacatctacgtattagtcatcgctattaccatggtcgaggtgagccccacgttctgcttcactctccccatctcccccccctccccacccccaattttgtatttatttattttttaattattttgtgcagcgatgggggcggggggggggggggggcgcgcgccaggcggggcggggcggggcgaggggcggggcggggcgaggcggagaggtgcggcggcagccaatcagagcggcgcgctccgaaagtttccttttatggcgaggcggcggcggcggcggccctataaaaagcgaagcgcgcggcgggcgggagtcgctgcgacgctgccttcgccccgtgccccgctccgccgccgcctcgcgccgcccgccccggctctgactgaccgcgttactcccacaggtgagcgggcgggacggcccttctcctccgggctgtaattagctgagcaagaggtaagggtttaagggatggttggttggtggggtattaatgtttaattacctggagcacctgcctgaaatcactttttttcaggttggaccggtgccaccatg*gactataaggaccacgacggagactacaaggatcatgatattgattacaaagacgatgacgataag*atggccccaaagaagaagcggaaggtcggtatccacggagtcccagcagccgacaagaagtacagcatcggcctggacatcggcaccaactctgtgggctgggccgtgatcaccgacgagtacaaggtgcccagcaagaaattcaaggtgctgggcaacaccgaccggcacagcatcaagaagaacctgatcggagccctgctgttcgacagcggcgaaacagccgaggccacccggctgaagagaaccgccagaagaagatacaccagacggaagaaccggatctgctatctgcaagagatcttcagcaacgagatggccaaggtggacgacagcttcttccacagactggaagagtccttcctggtggaagaggataagaagcacgagcggcaccccatcttcggcaacatcgtggacgaggtggcctaccacgagaagtaccccaccatctaccacctgagaaagaaactggtggacagcaccgacaaggccgacctgcggctgatctatctggccctggcccacatgatcaagttccggggccacttcctgatcgagggcgacctgaaccccgacaacagcgacgtggacaagctgttcatccagctggtgcagacctacaaccagctgttcgaggaaaaccccatcaacgccagcggcgtggacgccaaggccatcctgtctgccagactgagcaagagcagacggctggaaaatctgatcgcccagctgcccggcgagaagaagaatggcctgttcggaaacctgattgccctgagcctgggcctgacccccaacttcaagagcaacttcgacctggccgaggatgccaaactgcagctgagcaaggacacctacgacgacgacctggacaacctgctggcccagatcggcgaccagtacgccgacctgtttctggccgccaagaacctgtccgacgccatcctgctgagcgacatcctgagagtgaacaccgagatcaccaaggcccccctgagcgcctctatgatcaagagatacgacgagcaccaccaggacctgaccctgctgaaagctctcgtgcggcagcagctgcctgagaagtacaaagagattttcttcgaccagagcaagaacggctacgccggctacattgacggcggagccagccaggaagagttctacaagttcatcaagcccatcctggaaaagatggacggcaccgaggaactgctcgtgaagctgaacagagaggacctgctgcggaagcagcggaccttcgacaacggcagcatcccccaccagatccacctgggagagctgcacgccattctgcggcggcaggaagatttttacccattcctgaaggacaaccgggaaaagatcgagaagatcctgaccttccgcatcccctactacgtgggccctctggccaggggaaacagcagattcgcctggatgaccagaaagagcgaggaaaccatcaccccctggaacttcgaggaagtggtggacaagggcgcttccgcccagagcttcatcgagcggatgaccaacttcgataagaacctgcccaacgagaaggtgctgcccaagcacagcctgctgtacgagtacttcaccgtgtataacgagctgaccaaagtgaaatacgtgaccgagggaatgagaaagcccgccttcctgagcggcgagcagaaaaaggccatcgtggacctgctgttcaagaccaaccggaaagtgaccgtgaagcagctgaaagaggactacttcaagaaaatcgagtgcttcgactccgtggaaatctccggcgtggaagatcggttcaacgcctccctgggcacataccacgatctgctgaaaattatcaaggacaaggacttcctggacaatgaggaaaacgaggacattctggaagatatcgtgctgaccctgacactgtttgaggacagagagatgatcgaggaacggctgaaaacctatgcccacctgttcgacgacaaagtgatgaagcagctgaagcggcggagatacaccggctggggcaggctgagccggaagctgatcaacggcatccgggacaagcagtccggcaagacaatcctggatttcctgaagtccgacggcttcgccaacagaaacttcatgcagctgatccacgacgacagcctgacctttaaagaggacatccagaaagcccaggtgtccggccagggcgatagcctgcacgagcacattgccaatctggccggcagccccgccattaagaagggcatcctgcagacagtgaaggtggtggacgagctcgtgaaagtgatgggccggcacaagcccgagaacatcgtgatcgaaatggccagagagaaccagaccacccagaagggacagaagaacagccgcgagagaatgaagcggatcgaagagggcatcaaagagctgggcagccagatcctgaaagaacaccccgtggaaaacacccagctgcagaacgagaagctgtacctgtactacctgcagaatgggcgggatatgtacgtggaccaggaactggacatcaaccggctgtccgactacgatgtggaccatatcgtgcctcagagctttctgaaggacgactccatcgacaacaaggtgctgaccagaagcgacaagaaccggggcaagagcgacaacgtgccctccgaagaggtcgtgaagaagatgaagaactactggcggcagctgctgaacgccaagctgattacccagagaaagttcgacaatctgaccaaggccgagagaggcggcctgagcgaactggataaggccggcttcatcaagagacagctggtggaaacccggcagatcacaaagcacgtggcacagatcctggactcccggatgaacactaagtacgacgagaatgacaagctgatccgggaagtgaaagtgatcaccctgaagtccaagctggtgtccgatttCCGGAAGGATTTCCAGTTTTACAAAGTGCGCGAGATCAACAACTACCACCACGCCCACGACGCGTACCTGAACGCCGTCGTGGGAACCGCCCTGATCAAAAAGTACCCTaaGCTGGAAAGCGAGTTCGTGTACGGCGACTACAAGGTGTACGACGTaCGGAAGAtgatcgccaagagcgagcaggaaatcggcaaggctaccgccaagtacttcttctacagcaacatcatgaactttttcaagaccgagattaccctggccaacggcgagatccggaagcggcctctgatcgagacaaacggcgaaaccggggagatcgtgtgggataagggccgggattttgccaccgtgcggaaagtgctgagcatgccccaagtgaatatcgtgaaaaagaccgaggtgcagacaggcggcttcagcaaagagtctatcctgcccaagaggaacagcgataagctgatcgccagaaagaaggactgggaccctaagaagtacggcggcttcgacagccccaccgtggcctattctgtgctggtggtggccaaagtggaaaagggcaagtccaagaaactgaagagtgtgaaagagctgctggggatcaccatcatggaaagaagcagcttcgagaagaatcccatcgactttctggaagccaagggctacaaagaagtgaaaaaggacctgatcatcaagctgcctaagtactccctgttcgagctggaaaacggccggaagagaatgctggcctctgccggcgaactgcagaagggaaacgaactggccctgccctccaaatatgtgaacttcctgtacctggccagccactatgagaagctgaagggctcccccgaggataatgagcagaaacagctgtttgtggaacagcacaagcactacctggacgagatcatcgagcagatcagcgagttctccaagagagtgatcctggccgacgctaatctggacaaagtgctgtccgcctacaacaagcaccgggataagcccatcagagagcaggccgagaatatcatccacctgtttaccctgaccaatctgggagcccctgccgccttcaagtactttgacaccaccatcgaccggaagaggtacaccagcaccaaagaggtgctggacgccaccctgatccaccagagcatcaccggcctgtacgagacacggatcgacctgtctcagctgggaggcgacaaaaggccggcggccacgaaaaaggccggccaggcaaaaaagaaaaagtaagaattcctagagctcgctgatcagcctcgactgtgccttctagttgccagccatctgttgtttgcccctcccccgtgccttccttgaccctggaaggtgccactcccactgtcctttcctaataaaatgaggaaattgcatcgcattgtctgagtaggtgtcattctattctggggggtggggtggggcaggacagcaagggggaggattgggaagagaatagcaggcatgctggggagcggccgcaggaacccctagtgatggagttggccactccctctctgcgcgctcgctcgctcactgaggccgggcgaccaaaggtcgcccgacgcccgggctttgcccgggcggcctcagtgagcgagcgagcgcgcagctgcctgcaggggcgcctgatgcggtattttctccttacgcatctgtgcggtatttcacaccgcatacgtcaaagcaaccatagtacgcgccctgtagcggcgcattaagcgcggcgggtgtggtggttacgcgcagcgtgaccgctacacttgccagcgccctagcgcccgctcctttcgctttcttcccttcctttctcgccacgttcgccggctttccccgtcaagctctaaatcgggggctccctttagggttccgatttagtgctttacggcacctcgaccccaaaaaacttgatttgggtgatggttcacgtagtgggccatcgccctgatagacggtttttcgccctttgacgttggagtccacgttctttaatagtggactcttgttccaaactggaacaacactcaaccctatctcgggctattcttttgatttataagggattttgccgatttcggcctattggttaaaaaatgagctgatttaacaaaaatttaacgcgaattttaacaaaatattaacgtttacaattttatggtgcactctcagtacaatctgctctgatgccgcatagttaagccagccccgacacccgccaacacccgctgacgcgccctgacgggcttgtctgctcccggcatccgcttacagacaagctgtgaccgtctccgggagctgcatgtgtcagaggttttcaccgtcatcaccgaaacgcgcgagacgaaagggcctcgtgatacgcctatttttataggttaatgtcatgataataatggtttcttagacgtcaggtggcacttttcggggaaatgtgcgcggaacccctatttgtttatttttctaaatacattcaaatatgtatccgctcatgagacaataaccctgataaatgcttcaataatattgaaaaaggaagagtatgagtattcaacatttccgtgtcgcccttattcccttttttgcggcattttgccttcctgtttttgctcacccagaaacgctggtgaaagtaaaagatgctgaagatcagttgggtgcacgagtgggttacatcgaactggatctcaacagcggtaagatccttgagagttttcgccccgaagaacgttttccaatgatgagcacttttaaagttctgctatgtggcgcggtattatcccgtattgacgccgggcaagagcaactcggtcgccgcatacactattctcagaatgacttggttgagtactcaccagtcacagaaaagcatcttacggatggcatgacagtaagagaattatgcagtgctgccataaccatgagtgataacactgcggccaacttacttctgacaacgatcggaggaccgaaggagctaaccgcttttttgcacaacatgggggatcatgtaactcgccttgatcgttgggaaccggagctgaatgaagccataccaaacgacgagcgtgacaccacgatgcctgtagcaatggcaacaacgttgcgcaaactattaactggcgaactacttactctagcttcccggcaacaattaatagactggatggaggcggataaagttgcaggaccacttctgcgctcggcccttccggctggctggtttattgctgataaatctggagccggtgagcgtggaagccgcggtatcattgcagcactggggccagatggtaagccctcccgtatcgtagttatctacacgacggggagtcaggcaactatggatgaacgaaatagacagatcgctgagataggtgcctcactgattaagcattggtaactgtcagaccaagtttactcatatatactttagattgatttaaaacttcatttttaatttaaaaggatctaggtgaagatcctttttgataatctcatgaccaaaatcccttaacgtgagttttcgttccactgagcgtcagaccccgtagaaaagatcaaaggatcttcttgagatcctttttttctgcgcgtaatctgctgcttgcaaacaaaaaaaccaccgctaccagcggtggtttgtttgccggatcaagagctaccaactctttttccgaaggtaactggcttcagcagagcgcagataccaaatactgtccttctagtgtagccgtagttaggccaccacttcaagaactctgtagcaccgcctacatacctcgctctgctaatcctgttaccagtggctgctgccagtggcgataagtcgtgtcttaccgggttggactcaagacgatagttaccggataaggcgcagcggtcgggctgaacggggggttcgtgcacacagcccagcttggagcgaacgacctacaccgaactgagatacctacagcgtgagctatgagaaagcgccacgcttcccgaagggagaaaggcggacaggtatccggtaagcggcagggtcggaacaggagagcgcacgagggagcttccagggggaaacgcctggtatctttatagtcctgtcgggtttcgccacctctgacttgagcgtcgatttttgtgatgctcgtcaggggggcggagcctatggaaaaacgccagcaacgcggcctttttacggttcctggccttttgctggccttttgctcacatg

##### pX330-Flag-SpCas9-HF1 (Addgene #126755)

DNA sequence of the SpCas9-HF1 coding cassette:

Human codon optimized *S. pyogenes* Cas9 coloured in purple, NLS underlined, 3xFLAG tag in *light blue*, silent mutations underlined and marked with yellow*.*

atg*gactataaggaccacgacggagactacaaggatcatgatattgattacaaagacgatgacgataag*atggccccaaagaagaagcggaaggtcggtatccacggagtcccagcagccGACAAGAAGTACAGCATCGGCCTGGACATCGGCACCAACTCTGTGGGCTGGGCCGTGATCACCGACGAGTACAAGGTGCCCAGCAAGAAATTCAAGGTGCTGGGCAACACCGACCGGCACAGCATCAAGAAGAACCTGATCGGAGCCCTGCTGTTCGACAGCGGCGAAACAGCCGAGGCCACCCGGCTGAAGAGAACCGCCAGAAGAAGATACACCAGACGGAAGAACCGGATCTGCTATCTGCAAGAGATCTTCAGCAACGAGATGGCCAAGGTGGACGACAGCTTCTTCCACAGACTGGAAGAGTCCTTCCTGGTGGAAGAGGATAAGAAGCACGAGCGGCACCCCATCTTCGGCAACATCGTGGACGAGGTGGCCTACCACGAGAAGTACCCCACCATCTACCACCTGAGAAAGAAACTGGTGGACAGCACCGACAAGGCCGACCTGCGGCTGATCTATCTGGCCCTGGCCCACATGATCAAGTTCCGGGGCCACTTCCTGATCGAGGGCGACCTGAACCCCGACAACAGCGACGTGGACAAGCTGTTCATCCAGCTGGTGCAGACCTACAACCAGCTGTTCGAGGAAAACCCCATCAACGCCAGCGGCGTGGACGCCAAGGCCATCCTGTCTGCCAGACTGAGCAAGAGCAGACGGCTGGAAAATCTGATCGCCCAGCTGCCCGGCGAGAAGAAGAATGGCCTGTTCGGAAACCTGATTGCCCTGAGCCTGGGCCTGACCCCCAACTTCAAGAGCAACTTCGACCTGGCCGAGGATGCCAAACTGCAGCTGAGCAAGGACACCTACGACGACGACCTGGACAACCTGCTGGCCCAGATCGGCGACCAGTACGCCGACCTGTTTCTGGCCGCCAAGAACCTGTCCGACGCCATCCTGCTGAGCGACATCCTGAGAGTGAACACCGAGATCACCAAGGCCCCCCTGAGCGCCTCTATGATCAAGAGATACGACGAGCACCACCAGGACCTGACCCTGCTGAAAGCTCTCGTGCGGCAGCAGCTGCCTGAGAAGTACAAAGAGATTTTCTTCGACCAGAGCAAGAACGGCTACGCCGGCTACATTGACGGCGGAGCCAGCCAGGAAGAGTTCTACAAGTTCATCAAGCCCATCCTGGAAAAGATGGACGGCACCGAGGAACTGCTCGTGAAGCTGAACAGAGAGGACCTGCTGCGGAAGCAGCGGACCTTCGACAACGGCAGCATCCCCCACCAGATCCACCTGGGAGAGCTGCACGCCATTCTGCGGCGGCAGGAAGATTTTTACCCATTCCTGAAGGACAACCGGGAAAAGATCGAGAAGATCCTGACCTTCCGCATCCCCTACTACGTGGGCCCTCTGGCCAGGGGAAACAGCAGATTCGCCTGGATGACCAGAAAGAGCGAGGAAACCATCACCCCCTGGAACTTCGAGGAAGTGGTGGACAAGGGCGCTTCCGCCCAGAGCTTCATCGAGCGGATGACCgccTTCGATAAGAACCTGCCCAACGAGAAGGTGCTGCCCAAGCACAGCCTGCTGTACGAGTACTTCACCGTGTATAACGAGCTGACCAAAGTGAAATACGTGACCGAGGGAATGAGAAAGCCCGCCTTCCTGAGCGGCGAGCAGAAAAAGGCCATCGTGGACCTGCTGTTCAAGACCAACCGGAAAGTGACCGTGAAGCAGCTGAAAGAGGACTACTTCAAGAAAATCGAGTGCTTCGACTCCGTGGAAATCTCCGGCGTGGAAGATCGGTTCAACGCCTCCCTGGGCACATACCACGATCTGCTGAAAATTATCAAGGACAAGGACTTCCTGGACAATGAGGAAAACGAGGACATTCTGGAAGATATCGTGCTGACCCTGACACTGTTTGAGGACAGAGAGATGATCGAGGAACGGCTGAAAACCTATGCCCACCTGTTCGACGACAAAGTGATGAAGCAGCTGAAGCGGCGGAGATACACCGGCTGGGGCgcGCTGAGCCGGAAGCTGATCAACGGCATCCGcGACAAGCAGTCCGGCAAGACAATCCTGGATTTCCTGAAGTCCGACGGCTTCGCCAACAGAAACTTCATGgcGCTGATCCACGACGACAGCCTGACCTTTAAAGAGGACATCCAGAAAGCCCAGGTGTCCGGCCAGGGCGATAGCCTGCACGAGCACATTGCCAATCTcGCCGGCAGCCCCGCCATTAAGAAGGGCATCCTcCAGACAGTGAAGGTGGTGGACGAGCTgGTGAAAGTGATGGGCCGGCACAAGCCCGAGAACATCGTGATCGAAATGGCCAGAGAGAACCAGACCACCCAGAAGGGACAGAAGAACAGtCGCGAGAGAATGAAGCGGATCGAAGAGGGCATCAAAGAGCTGGGCAGCCAGATCCTGAAAGAACACCCCGTGGAAAACACCCAGCTcCAGAACGAGAAGCTGTACCTGTACTACCTcCAGAATGGGCGGGATATGTACGTGGACCAGGAACTGGACATCAACCGGCTGTCCGACTACGATGTGGACCATATCGTGCCTCAGAGCTTTCTtaaGGACGACTCCATCGACAACAAGGTGCTGACCAGAAGCGACAAGAACCGGGGCAAGAGCGACAACGTGCCCTCCGAAGAGGTCGTGAAGAAGATGAAGAACTACTGGCGGCAGCTGCTGAACGCCAAGCTGATTACCCAGAGAAAGTTCGACAATCTGACCAAGGCCGAGAGAGGCGGCCTGAGCGAACTGGATAAGGCCGGCTTCATCAAGAGACAGCTGGTGGAAACCCGGgcGatcacaaagcacgtggcacagatcctggactcccggatgaacactaagtacgacgagaatgacaagctgatccgggaagtgaaagtgatcaccctgaagtccaagctggtgtccgatttCCGGAAGGATTTCCAGTTTTACAAAGTGCGCGAGATCAACAACTACCACCACGCCCACGACGCGTACCTGAACGCCGTCGTGGGAACCGCCCTGATCAAAAAGTACCCTaaGCTGGAAAGCGAGTTCGTGTACGGCGACTACAAGGTGTACGACGTaCGGAAGAtgatcgccaagagcgagcaggaaatcggcaaggctaccgccaagtacttcttctacagcaacatcatgaactttttcaagaccgagattaccctggccaacggcgagatccggaagcggcctctgatcgagacaaacggcgaaaccggggagatcgtgtgggataagggccgggattttgccaccgtgcggaaagtgctgagcatgccccaagtgaatatcgtgaaaaagaccgaggtgcagacaggcggcttcagcaaagagtctatcctgcccaagaggaacagcgataagctgatcgccagaaagaaggactgggaTcctaagaagtacggcggcttcgacagccccaccgtggcctattctgtgctggtggtggccaaagtggaaaagggcaagtccaagaaGctTaagagtgtgaaagagctgctggggatcaccatcatggaaagaagcagcttcgagaagaatcccatcgactttctggaagccaagggctacaaagaagtgaaaaaggacctgatcatcaagctgcctaagtactcGctCttcgagctggaaaacggccggaagagaatgctggcctctgccggcgaactgcagaagggaaacgaactggccctgccctccaaatatgtgaacttcctgtacctggccagccactatgagaagctgaagggctcccccgaggataatgagcagaaacagctgtttgtggaacagcacaagcactacctggacgagatcatcgagcagatcagcgagttctccaagagagtgatcctggccgacgctaatctggacaaagtgctgtccgcctacaacaagcaccgggataagcccatcagagagcaggccgagaatatcatccacctgtttaccctgaccaatctgggagcccctgccgccttcaagtactttgacaccaccatcgaccggaagaggtacaccagcaccaaagaggtgctggacgccaccctgatccaccagagcatcaccggcctgtacgagacacggatcgacctgtctcagctgggaggcgacAAAAGGCCGGCGGCCACGAAAAAGGCCGGccaggcaaaaaagaaaaagtaa

##### pX330-Flag-HypaSpCas9 (Addgene #126756)

DNA sequence of the HypaSpCas9 coding cassette:

Human codon optimized *S. pyogenes* Cas9 coloured in purple, NLS underlined, 3xFLAG tag in *light blue*, silent mutations underlined and marked with yellow*.*

atg*gactataaggaccacgacggagactacaaggatcatgatattgattacaaagacgatgacgataag*atggccccaaagaagaagcggaaggtcggtatccacggagtcccagcagccgaCAAGAAGTACAGCATCGGCCTGGACATCGGCACCAACTCTGTGGGCTGGGCCGTGATCACCGACGAGTACAAGGTGCCCAGCAAGAAATTCAAGGTGCTGGGCAACACCGACCGGCACAGCATCAAGAAGAACCTGATCGGAGCCCTGCTGTTCGACAGCGGCGAAACAGCCGAGGCCACCCGGCTGAAGAGAACCGCCAGAAGAAGATACACCAGACGGAAGAACCGGATCTGCTATCTGCAAGAGATCTTCAGCAACGAGATGGCCAAGGTGGACGACAGCTTCTTCCACAGACTGGAAGAGTCCTTCCTGGTGGAAGAGGATAAGAAGCACGAGCGGCACCCCATCTTCGGCAACATCGTGGACGAGGTGGCCTACCACGAGAAGTACCCCACCATCTACCACCTGAGAAAGAAACTGGTGGACAGCACCGACAAGGCCGACCTGCGGCTGATCTATCTGGCCCTGGCCCACATGATCAAGTTCCGGGGCCACTTCCTGATCGAGGGCGACCTGAACCCCGACAACAGCGACGTGGACAAGCTGTTCATCCAGCTGGTGCAGACCTACAACCAGCTGTTCGAGGAAAACCCCATCAACGCCAGCGGCGTGGACGCCAAGGCCATCCTGTCTGCCAGACTGAGCAAGAGCAGACGGCTGGAAAATCTGATCGCCCAGCTGCCCGGCGAGAAGAAGAATGGCCTGTTCGGAAACCTGATTGCCCTGAGCCTGGGCCTGACCCCCAACTTCAAGAGCAACTTCGACCTGGCCGAGGATGCCAAACTGCAGCTGAGCAAGGACACCTACGACGACGACCTGGACAACCTGCTGGCCCAGATCGGCGACCAGTACGCCGACCTGTTTCTGGCCGCCAAGAACCTGTCCGACGCCATCCTGCTGAGCGACATCCTGAGAGTGAACACCGAGATCACCAAGGCCCCCCTGAGCGCCTCTATGATCAAGAGATACGACGAGCACCACCAGGACCTGACCCTGCTGAAAGCTCTCGTGCGGCAGCAGCTGCCTGAGAAGTACAAAGAGATTTTCTTCGACCAGAGCAAGAACGGCTACGCCGGCTACATTGACGGCGGAGCCAGCCAGGAAGAGTTCTACAAGTTCATCAAGCCCATCCTGGAAAAGATGGACGGCACCGAGGAACTGCTCGTGAAGCTGAACAGAGAGGACCTGCTGCGGAAGCAGCGGACCTTCGACAACGGCAGCATCCCCCACCAGATCCACCTGGGAGAGCTGCACGCCATTCTGCGGCGGCAGGAAGATTTTTACCCATTCCTGAAGGACAACCGGGAAAAGATCGAGAAGATCCTGACCTTCCGCATCCCCTACTACGTGGGCCCTCTGGCCAGGGGAAACAGCAGATTCGCCTGGATGACCAGAAAGAGCGAGGAAACCATCACCCCCTGGAACTTCGAGGAAGTGGTGGACAAGGGCGCTTCCGCCCAGAGCTTCATCGAGCGGATGACCaaCTTCGATAAGAACCTGCCCAACGAGAAGGTGCTGCCCAAGCACAGCCTGCTGTACGAGTACTTCACCGTGTATAACGAGCTGACCAAAGTGAAATACGTGACCGAGGGAATGAGAAAGCCCGCCTTCCTGAGCGGCGAGCAGAAAAAGGCCATCGTGGACCTGCTGTTCAAGACCAACCGGAAAGTGACCGTGAAGCAGCTGAAAGAGGACTACTTCAAGAAAATCGAGTGCTTCGACTCCGTGGAAATCTCCGGCGTGGAAGATCGGTTCAACGCCTCCCTGGGCACATACCACGATCTGCTGAAAATTATCAAGGACAAGGACTTCCTGGACAATGAGGAAAACGAGGACATTCTGGAAGATATCGTGCTGACCCTGACACTGTTTGAGGACAGAGAGATGATCGAGGAACGGCTGAAAACCTATGCCCACCTGTTCGACGACAAAGTGATGAAGCAGCTGAAGCGGCGGAGATACACCGGCTGGGGCAGGCTGAGCCGGAAGCTGATCAACGGCATCCGcGACAAGCAGTCCGGCAAGACAATCCTGGATTTCCTGAAGTCCGACGGCTTCGCCAACAGAGCCTTtgcagccCTGATCgctgacgacagcctgacctttaaagaggacatccagaaagcccaggtgtccggccagggcgatagcctgcacgagcacattgccaatctggccggcagccccgccattaagaagggcatcctgcagacagtgaaggtggtggacgagctcgtgaaagtgatgggccggcacaagcccgagaacatcgtgatcgaaatggccagagagaaccagaccacccagaagggacagaagaacagccgcgagagaatgaagcggatcgaagagggcatcaaagagctgggcagccagatcctgaaagaacaccccgtggaaaacacccagctgcagaacgagaagctgtacctgtactacctgcagaatgggcgggatatgtacgtggaccaggaactggacatcaaccggctgtccgactacgatgtggaccatatcgtgcctcagagctttctgaaggacgactccatcgacaacaaggtgctgaccagaagcgacaagaaccggggcaagagcgacaacgtgccctccgaagaggtcgtgaagaagatgaagaactactggcggcagctgctgaacgccaagctgattacccagagaaagttcgacaatctgaccaaggccgagagaggcggcctgagcgaactggataaggccggcttcatcaagagacagctggtggaaacccggcagatcacaaagcacgtggcacagatcctggactcccggatgaacactaagtacgacgagaatgacaagctgatccgggaagtgaaagtgatcaccctgaagtccaagctggtgtccgatttCCGGAAGGATTTCCAGTTTTACAAAGTGCGCGAGATCAACAACTACCACCACGCCCACGACGCGTACCTGAACGCCGTCGTGGGAACCGCCCTGATCAAAAAGTACCCTaaGCTGGAAAGCGAGTTCGTGTACGGCGACTACAAGGTGTACGACGTaCGGAAGATGATCGCCAAGAGCGAGCAGGAAATCGGCAAGGCTACCGCCAAGTACTTCTTCTACAGCAACATCATGAACTTTTTCAAGACCGAGATTACCCTGGCCAACGGCGAGATccggaagcggcctctgatcgagacaaacggcgaaaccggggagatcgtgtgggataagggccgggattttgccaccgtgcggaaagtgctgagcatgccccaagtgaatatcgtgaaaaagaccgaggtgcagacaggcggcttcagcaaagagtctatcctgcccaagaggaacagcgataagctgatcgccagaaagaaggactgggaccctaagaagtacggcggcttcgacagccccaccgtggcctattctgtgctggtggtggccaaagtggaaaagggcaagtccaagaaactgaagagtgtgaaagagctgctggggatcaccatcatggaaagaagcagcttcgagaagaatcccatcgactttctggaagccaagggctacaaagaagtgaaaaaggacctgatcatcaagctgcctaagtactccctgttcgagctggaaaacggccggaagagaatgctggcctctgccggcgaactgcagaagggaaacgaactggccctgccctccaaatatgtgaacttcctgtacctggccagccactatgagaagctgaagggctcccccgaggataatgagcagaaacagctgtttgtggaacagcacaagcactacctggacgagatcatcgagcagatcagcgagttctccaagagagtgatcctggccgacgctaatctggacaaagtgctgtccgcctacaacaagcaccgggataagcccatcagagagcaggccgagaatatcatccacctgtttaccctgaccaatctgggagcccctgccgccttcaagtactttgacaccaccatcgaccggaagaggtacaccagcaccaaagaggtgctggacgccaccctgatccaccagagcatcaccggcctgtacgagacacggatcgacctgtctcagctgggaggcgacAAAAGGCCGGCGGCCACGAAAAAGGCCGGCCAGGCAAAAAAGAAAAAGtaa

##### pX330-Flag-evoSpCas9 (Addgene #126758)

DNA sequence of the HypaSpCas9 coding cassette:

Human codon optimized *S. pyogenes* Cas9 coloured in purple, NLS underlined, 3xFLAG tag in *light blue*, silent mutations underlined and marked with yellow*.*

atg*gactataaggaccacgacggagactacaaggatcatgatattgattacaaagacgatgacgataag*atggccccaaagaagaagcggaaggtcggtatccacggagtcccagcagccgaCAAGAAGTACAGCATCGGCCTGGACATCGGCACCAACTCTGTGGGCTGGGCCGTGATCACCGACGAGTACAAGGTGCCCAGCAAGAAATTCAAGGTGCTGGGCAACACCGACCGGCACAGCATCAAGAAGAACCTGATCGGAGCCCTGCTGTTCGACAGCGGCGAAACAGCCGAGGCCACCCGGCTGAAGAGAACCGCCAGAAGAAGATACACCAGACGGAAGAACCGGATCTGCTATCTGCAAGAGATCTTCAGCAACGAGATGGCCAAGGTGGACGACAGCTTCTTCCACAGACTGGAAGAGTCCTTCCTGGTGGAAGAGGATAAGAAGCACGAGCGGCACCCCATCTTCGGCAACATCGTGGACGAGGTGGCCTACCACGAGAAGTACCCCACCATCTACCACCTGAGAAAGAAACTGGTGGACAGCACCGACAAGGCCGACCTGCGGCTGATCTATCTGGCCCTGGCCCACATGATCAAGTTCCGGGGCCACTTCCTGATCGAGGGCGACCTGAACCCCGACAACAGCGACGTGGACAAGCTGTTCATCCAGCTGGTGCAGACCTACAACCAGCTGTTCGAGGAAAACCCCATCAACGCCAGCGGCGTGGACGCCAAGGCCATCCTGTCTGCCAGACTGAGCAAGAGCAGACGGCTGGAAAATCTGATCGCCCAGCTGCCCGGCGAGAAGAAGAATGGCCTGTTCGGAAACCTGATTGCCCTGAGCCTGGGCCTGACCCCCAACTTCAAGAGCAACTTCGACCTGGCCGAGGATGCCAAACTGCAGCTGAGCAAGGACACCTACGACGACGACCTGGACAACCTGCTGGCCCAGATCGGCGACCAGTACGCCGACCTGTTTCTGGCCGCCAAGAACCTGTCCGACGCCATCCTGCTGAGCGACATCCTGAGAGTGAACACCGAGATCACCAAGGCCCCCCTGAGCGCCTCTATGATCAAGAGATACGACGAGCACCACCAGGACCTGACCCTGCTGAAAGCTCTCGTGCGGCAGCAGCTGCCTGAGAAGTACAAAGAGATTTTCTTCGACCAGAGCAAGAACGGCTACGCCGGCTACATTGACGGCGGAGCCAGCCAGGAAGAGTTCTACAAGTTCATCAAGCCCATCCTGGAAAAGATGGACGGCACCGAGGAACTGCTCGTGAAGCTGAACAGAGAGGACCTGCTGCGGAAGCAGCGGACCTTCGACAACGGCAGCATCCCCCACCAGATCCACCTGGGAGAGCTGCACGCCATTCTGCGGCGGCAGGAAGATTTTTACCCATTCCTGAAGGACAACCGGGAAAAGATCGAGAAGATCCTGACCTTCCGCATCCCCTACTACGTGGGCCCTCTGGCCAGGGGAAACAGCAGATTCGCCTGGATGACCAGAAAGAGCGAGGAAACCATCACCCCCTGGAACTTCGAGGAAGTGGTGGACAAGGGCGCTTCCGCCCAGAGCTTCATCGAGCGGgtgACCaaCTTCGATAAGAACCTGCCCAACGAGAAGGTGCTGCCCAAGCACAGCCTGCTGaACGAGTACTTCACCGTGTATAACGAGCTGACCgAAGTGAAATACGTGACCGAGGGAATGAGAAAGCCCGCCTTCCTGAGCGGCGAGCAGAAAAAGGCCATCGTGGACCTGCTGTTCAAGACCAACCGGAAAGTGACCGTGAAGCAGCTGAAAGAGGACTACTTCAAGAAAATCGAGTGCTTCGACTCCGTGGAAATCTCCGGCGTGGAAGATCGGTTCAACGCCTCCCTGGGCACATACCACGATCTGCTGAAAATTATCAAGGACAAGGACTTCCTGGACAATGAGGAAAACGAGGACATTCTGGAAGATATCGTGCTGACCCTGACACTGTTTGAGGACAGAGAGATGATCGAGGAACGGCTGAAAACCTATGCCCACCTGTTCGACGACAAAGTGATGAAGCAGCTGAAGCGGCGGAGATACACCGGCTGGGGCcaGCTGAGCCGGAAGCTGATCAACGGCATCCGgGACAAGCAGTCCGGCAAGACAATCCTGGATTTCCTGAAGTCCGACGGCTTCGCCAACAGAaacttcatgcagctgatccacgacgacagcctgacctttaaagaggacatccagaaagcccaggtgtccggccagggcgatagcctgcacgagcacattgccaatctggccggcagccccgccattaagaagggcatcctgcagacagtgaaggtggtggacgagctcgtgaaagtgatgggccggcacaagcccgagaacatcgtgatcgaaatggccagagagaaccagaccacccagaagggacagaagaacagccgcgagagaatgaagcggatcgaagagggcatcaaagagctgggcagccagatcctgaaagaacaccccgtggaaaacacccagctgcagaacgagaagctgtacctgtactacctgcagaatgggcgggatatgtacgtggaccaggaactggacatcaaccggctgtccgactacgatgtggaccatatcgtgcctcagagctttctgaaggacgactccatcgacaacaaggtgctgaccagaagcgacaagaaccggggcaagagcgacaacgtgccctccgaagaggtcgtgaagaagatgaagaactactggcggcagctgctgaacgccaagctgattacccagagaaagttcgacaatctgaccaaggccgagagaggcggcctgagcgaactggataaggccggcttcatcaagagacagctggtggaaacccggcagatcacaaagcacgtggcacagatcctggactcccggatgaacactaagtacgacgagaatgacaagctgatccgggaagtgaaagtgatcaccctgaagtccaagctggtgtccgatttCCGGAAGGATTTCCAGTTTTACAAAGTGCGCGAGATCAACAACTACCACCACGCCCACGACGCGTACCTGAACGCCGTCGTGGGAACCGCCCTGATCAAAAAGTACCCTaaGCTGGAAAGCGAGTTCGTGTACGGCGACTACAAGGTGTACGACGTaCGGAAGATGATCGCCAAGAGCGAGCAGGAAATCGGCAAGGCTACCGCCAAGTACTTCTTCTACAGCAACATCATGAACTTTTTCAAGACCGAGATTACCCTGGCCAACGGCGAGATccggaagcggcctctgatcgagacaaacggcgaaaccggggagatcgtgtgggataagggccgggattttgccaccgtgcggaaagtgctgagcatgccccaagtgaatatcgtgaaaaagaccgaggtgcagacaggcggcttcagcaaagagtctatcctgcccaagaggaacagcgataagctgatcgccagaaagaaggactgggaccctaagaagtacggcggcttcgacagccccaccgtggcctattctgtgctggtggtggccaaagtggaaaagggcaagtccaagaaactgaagagtgtgaaagagctgctggggatcaccatcatggaaagaagcagcttcgagaagaatcccatcgactttctggaagccaagggctacaaagaagtgaaaaaggacctgatcatcaagctgcctaagtactccctgttcgagctggaaaacggccggaagagaatgctggcctctgccggcgaactgcagaagggaaacgaactggccctgccctccaaatatgtgaacttcctgtacctggccagccactatgagaagctgaagggctcccccgaggataatgagcagaaacagctgtttgtggaacagcacaagcactacctggacgagatcatcgagcagatcagcgagttctccaagagagtgatcctggccgacgctaatctggacaaagtgctgtccgcctacaacaagcaccgggataagcccatcagagagcaggccgagaatatcatccacctgtttaccctgaccaatctgggagcccctgccgccttcaagtactttgacaccaccatcgaccggaagaggtacaccagcaccaaagaggtgctggacgccaccctgatccaccagagcatcaccggcctgtacgagacacggatcgacctgtctcagctgggaggcgacAAAAGGCCGGCGGCCACGAAAAAGGCCGGCCAGGCAAAAAAGAAAAAGtaa

##### pPIK16045_pX330_Flag-SuperFi-Cas9 (Addgene #184370)

DNA sequence of the HiFiSpCas9 coding cassette:

Human codon optimized *S. pyogenes* Cas9 coloured in purple, NLS underlined, 3xFLAG tag in *light blue*, silent mutations underlined and marked with yellow*.*

atg*gactataaggaccacgacggagactacaaggatcatgatattgattacaaagacgatgacgataag*atggccccaaagaagaagcggaaggtcggtatccacggagtcccagcagccgacaagaagtacagcatcggcctggacatcggcaccaactctgtgggctgggccgtgatcaccgacgagtacaaggtgcccagcaagaaattcaaggtgctgggcaacaccgaccggcacagcatcaagaagaacctgatcggagccctgctgttcgacagcggcgaaacagccgaggccacccggctgaagagaaccgccagaagaagatacaccagacggaagaaccggatctgctatctgcaagagatcttcagcaacgagatggccaaggtggacgacagcttcttccacagactggaagagtccttcctggtggaagaggataagaagcacgagcggcaccccatcttcggcaacatcgtggacgaggtggcctaccacgagaagtaccccaccatctaccacctgagaaagaaactggtggacagcaccgacaaggccgacctgcggctgatctatctggccctggcccacatgatcaagttccggggccacttcctgatcgagggcgacctgaaccccgacaacagcgacgtggacaagctgttcatccagctggtgcagacctacaaccagctgttcgaggaaaaccccatcaacgccagcggcgtggacgccaaggccatcctgtctgccagactgagcaagagcagacggctggaaaatctgatcgcccagctgcccggcgagaagaagaatggcctgttcggaaacctgattgccctgagcctgggcctgacccccaacttcaagagcaacttcgacctggccgaggatgccaaactgcagctgagcaaggacacctacgacgacgacctggacaacctgctggcccagatcggcgaccagtacgccgacctgtttctggccgccaagaacctgtccgacgccatcctgctgagcgacatcctgagagtgaacaccgagatcaccaaggcccccctgagcgcctctatgatcaagagatacgacgagcaccaccaggacctgaccctgctgaaagctctcgtgcggcagcagctgcctgagaagtacaaagagattttcttcgaccagagcaagaacggctacgccggctacattgacggcggagccagccaggaagagttctacaagttcatcaagcccatcctggaaaagatggacggcaccgaggaactgctcgtgaagctgaacagagaggacctgctgcggaagcagcggaccttcgacaacggcagcatcccccaccagatccacctgggagagctgcacgccattctgcggcggcaggaagatttttacccattcctgaaggacaaccgggaaaagatcgagaagatcctgaccttccgcatcccctactacgtgggccctctggccaggggaaacagcagattcgcctggatgaccagaaagagcgaggaaaccatcaccccctggaacttcgaggaagtggtggacaagggcgcttccgcccagagcttcatcgagcggatgaccaacttcgataagaacctgcccaacgagaaggtgctgcccaagcacagcctgctgtacgagtacttcaccgtgtataacgagctgaccaaagtgaaatacgtgaccgagggaatgagaaagcccgccttcctgagcggcgagcagaaaaaggccatcgtggacctgctgttcaagaccaaccggaaagtgaccgtgaagcagctgaaagaggactacttcaagaaaatcgagtgcttcgactccgtggaaatctccggcgtggaagatcggttcaacgcctccctgggcacataccacgatctgctgaaaattatcaaggacaaggacttcctggacaatgaggaaaacgaggacattctggaagatatcgtgctgaccctgacactgtttgaggacagagagatgatcgaggaacggctgaaaacctatgcccacctgttcgacgacaaagtgatgaagcagctgaagcggcggagatacaccggctggggcaggctgagccggaagctgatcaacggcatccgggacaagcagtccggcaagacaatcctggatttcctgaagtccgacggcttcgccaacagaaacttcatgcagctgatccacgacgacagcctgacctttaaagaggacatccagaaagcccaggtgtccggccagggcgatagcctgcacgagcacattgccaatctggccggcagccccgccattaagaagggcatcctgcagacagtgaaggtggtggacgagctcgtgaaagtgatgggccggcacaagcccgagaacatcgtgatcgaaatggccagagagaaccagaccacccagaagggacagaagaacagccgcgagagaatgaagcggatcgaagagggcatcaaagagctgggcagccagatcctgaaagaacaccccgtggaaaacacccagctgcagaacgagaagctgtacctgtactacctgcagaatgggcgggatatgtacgtggaccaggaactggacatcaaccggctgtccgactacgatgtggaccatatcgtgcctcagagctttctgaaggacgactccatcgacaacaaggtgctgaccagaagcgacaagaaccggggcaagagcgacaacgtgccctccgaagaggtcgtgaagaagatgaagaactactggcggcagctgctgaacgccaagctgattacccagagaaagttcgacaatctgaccaaggccgagagaggcggcctgagcgaactggataaggccggcttcatcaagagacagctggtggaaacccggcagatcacaaagcacgtggcacagatcctggactcccggatgaacactaagtacgacgagaatgacaagctgatccgggaagtgaaagtgatcaccctgaagtccaagctggtgtccgatttCCGGAAGGATTTCCAGTTTTACAAAGTGCGCGAGATCAACAACTACCACCACGCCCACGACGCGTACCTGAACGCCGTCGTGGGAACCGCCCTGATCAAAAAGTACCCTaaGCTGGAAAGCGAGTTCGTGgacGGCGACgacAAGGTGgacGACgacgacAAGATGATCGCCAAGAGCGAagacGAAATCGGCgacgctaccgccaagtacttcttctacagcaacatcatgaactttttcaagaccgagattaccctggccaacggcgagatccggaagcggcctctgatcgagacaaacggcgaaaccggggagatcgtgtgggataagggccgggattttgccaccgtgcggaaagtgctgagcatgccccaagtgaatatcgtgaaaaagaccgaggtgcagacaggcggcttcagcaaagagtctatcctgcccaagaggaacagcgataagctgatcgccagaaagaaggactgggaccctaagaagtacggcggcttcgacagccccaccgtggcctattctgtgctggtggtggccaaagtggaaaagggcaagtccaagaaactgaagagtgtgaaagagctgctggggatcaccatcatggaaagaagcagcttcgagaagaatcccatcgactttctggaagccaagggctacaaagaagtgaaaaaggacctgatcatcaagctgcctaagtactccctgttcgagctggaaaacggccggaagagaatgctggcctctgccggcgaactgcagaagggaaacgaactggccctgccctccaaatatgtgaacttcctgtacctggccagccactatgagaagctgaagggctcccccgaggataatgagcagaaacagctgtttgtggaacagcacaagcactacctggacgagatcatcgagcagatcagcgagttctccaagagagtgatcctggccgacgctaatctggacaaagtgctgtccgcctacaacaagcaccgggataagcccatcagagagcaggccgagaatatcatccacctgtttaccctgaccaatctgggagcccctgccgccttcaagtactttgacaccaccatcgaccggaagaggtacaccagcaccaaagaggtgctggacgccaccctgatccaccagagcatcaccggcctgtacgagacacggatcgacctgtctcagctgggaggcgacAAAAGGCCGGCGGCCACGAAAAAGGCCGGCCAGGCAAAAAAGAAAAAGtaa

##### px330-Flag-B-HypaR-SpCas9 (Addgene #126764)

DNA sequence of the HypaR-SpCas9 coding cassette:

Human codon optimized *S. pyogenes* Cas9 shown in purple, NLS underlined, modified codons in red, 3xFLAG tag in *light blue*, silent mutations underlined and marked with yellow*,* insertions marked with green, deletions crossed out and marked with grey.

atg*gactataaggaccacgacggagactacaaggatcatgatattgattacaaagacgatgacgataag*atggccccaaagaagaagcggaaggtcggtatccacggagtcccagcagccgaCAAGAAGTACAGCATCGGCCTGGACATCGGCACCAACTCTGTGGGCTGGGCCGTGATCACCGACGAGTACAAGGTGCCCAGCAAGAAATTCAAGGTGCTGGGCAACACCGACCGGCACAGCATCAAGAAGAACCTGATCGGAGCCCTGCTGTTCGACAGCGGCGAAACAGCCGAGGCCACCCGGCTGAAGAGAACCGCCAGAAGAAGATACACCAGACGGAAGAACCGGATCTGCTATCTGCAAGAGATCTTCAGCAACGAGATGGCCAAGGTGGACGACAGCTTCTTCCACAGACTGGAAGAGTCCTTCCTGGTGGAAGAGGATAAGAAGCACGAGCGGCACCCCATCTTCGGCAACATCGTGGACGAGGTGGCCTACCACGAGAAGTACCCCACCATCTACCACCTGAGAAAGAAACTGGTGGACAGCACCGACAAGGCCGACCTGCGGCTGATCTATCTGGCCCTGGCCCACATGATCAAGTTCCGGGGCCACTTCCTGATCGAGGGCGACCTGAACCCCGACAACAGCGACGTGGACAAGCTGTTCATCCAGCTGGTGCAGACCTACAACCAGCTGTTCGAGGAAAACCCCATCAACGCCAGCGGCGTGGACGCCAAGGCCATCCTGTCTGCCAGACTGAGCAAGAGCAGACGGCTGGAAAATCTGATCGCCCAGCTGCCCGGCGAGAAGAAGAATGGCCTGTTCGGAAACCTGATTGCCCTGAGCCTGGGCCTGACCCCCAACTTCAAGAGCAACTTCGACCTGGCCGAGGATGCCAAACTGCAGCTGAGCAAGGACACCTACGACGACGACCTGGACAACCTGCTGGCCCAGATCGGCGACCAGTACGCCGACCTGTTTCTGGCCGCCAAGAACCTGTCCGACGCCATCCTGCTGAGCGACATCCTGAGAGTGAACACCGAGATCACCAAGGCCCCCCTGAGCGCCTCTATGATCAAGAGATACGACGAGCACCACCAGGACCTGACCCTGCTGAAAGCTCTCGTGCGGCAGCAGCTGCCTGAGAAGTACAAAGAGATTTTCTTCGACCAGAGCAAGAACGGCTACGCCGGCTACATTGACGGCGGAGCCAGCCAGGAAGAGTTCTACAAGTTCATCAAGCCCATCCTGGAAAAGATGGACGGCACCGAGGAACTGCTCGTGAAGCTGAACAGAGAGGACCTGCTGCGGAAGCAGCGGACCTTCGACAACGGCAGCATCCCCCACCAGATCCACCTGGGAGAGCTGCACGCCATTCTGCGGCGGCAGGAAGATTTTTACCCATTCCTGAAGGACAACCGGGAAAAGATCGAGAAGATCCTGACCTTCCGCATCCCCTACTACGTGGGCCCTCTGGCCAGGGGAAACAGCAGATTCGCCTGGATGACCAGAAAGAGCGAGGAAACCATCACCCCCTGGAACTTCGAGGAAGTGGTGGACAAGGGCGCTTCCGCCCAGAGCTTCATCGAGCGGATGACCaaCTTCGATAAGAACCTGCCCAACGAGAAGGTGCTGCCCAAGCACAGCCTGCTGTACGAGTACTTCACCGTGTATAACGAGCTGACCAAAGTGAAATACGTGACCGAGGGAATGAGAAAGCCCGCCTTCCTGAGCGGCGAGCAGAAAAAGGCCATCGTGGACCTGCTGTTCAAGACCAACCGGAAAGTGACCGTGAAGCAGCTGAAAGAGGACTACTTCAAGAAAATCGAGTGCTTCGACTCCGTGGAAATCTCCGGCGTGGAAGATCGGTTCAACGCCTCCCTGGGCACATACCACGATCTGCTGAAAATTATCAAGGACAAGGACTTCCTGGACAATGAGGAAAACGAGGACATTCTGGAAGATATCGTGCTGACCCTGACACTGTTTGAGGACAGAGAGATGATCGAGGAACGGCTGAAAACCTATGCCCACCTGTTCGACGACAAAGTGATGAAGCAGCTGAAGCGGCGGAGATACACCGGCTGGGGCGcGCTGAGCCGGAAGCTGATCAACGGCATCCGcGACAAGCAGTCCGGCAAGACAATCCTGGATTTCCTGAAGTCCGACGGCTTCGCCAACAGAGCCTTtgcagccCTGATCgctgacgacagcctgacctttaaagaggacatccagaaagcccaggtgtccggccagggcgatagcctgcacgagcacattgccaatctggccggcagccccgccattaagaagggcatcctgcagacagtgaaggtggtggacgagctcgtgaaagtgatgggccggcacaagcccgagaacatcgtgatcgaaatggccagagagaaccagaccacccagaagggacagaagaacagccgcgagagaatgaagcggatcgaagagggcatcaaagagctgggcagccagatcctgaaagaacaccccgtggaaaacacccagctgcagaacgagaagctgtacctgtactacctgcagaatgggcgggatatgtacgtggaccaggaactggacatcaaccggctgtccgactacgatgtggaccatatcgtgcctcagagctttctgaaggacgactccatcgacaacaaggtgctgaccagaagcgacaagaaccggggcaagagcgacaacgtgccctccgaagaggtcgtgaagaagatgaagaactactggcggcagctgctgaacgccaagctgattacccagagaaagttcgacaatctgaccaaggccgagagaggcggcctgagcgaactggataaggccggcttcatcaagagacagctggtggaaacccggcagatcacaaagcacgtggcacagatcctggactcccggatgaacactaagtacgacgagaatgacaagctgatccgggaagtgaaagtgatcaccctgaagtccaagctggtgtccgatttCCGGAAGGATTTCCAGTTTTACAAAGTGCGCGAGATCAACAACTACCACCACGCCCACGACGCGTACCTGAACGCCGTCGTGGGAACCGCCCTGATCAAAAAGTACCCTaaGCTGggtggc~~GAAAGCGAGTTCGTGTACGGCGACTAC~~AAGGTGTACGACGTaCGGAAGATGATCGCCAAGAGCGAGCAGGAAATCGGCAAGGCTACCGCCAAGTACTTCTTCTACAGCAACATCATGAACTTTTTCAAGACCGAGATTACCCTGGCCAACGGCGAGATccggaagcggcctctgatcgagacaaacggcgaaaccggggagatcgtgtgggataagggccgggattttgccaccgtgcggaaagtgctgagcatgccccaagtgaatatcgtgaaaaagaccgaggtgcagacaggcggcttcagcaaagagtctatcctgcccaagaggaacagcgataagctgatcgccagaaagaaggactgggaccctaagaagtacggcggcttcgacagccccaccgtggcctattctgtgctggtggtggccaaagtggaaaagggcaagtccaagaaactgaagagtgtgaaagagctgctggggatcaccatcatggaaagaagcagcttcgagaagaatcccatcgactttctggaagccaagggctacaaagaagtgaaaaaggacctgatcatcaagctgcctaagtactccctgttcgagctggaaaacggccggaagagaatgctggcctctgccggcgaactgcagaagggaaacgaactggccctgccctccaaatatgtgaacttcctgtacctggccagccactatgagaagctgaagggctcccccgaggataatgagcagaaacagctgtttgtggaacagcacaagcactacctggacgagatcatcgagcagatcagcgagttctccaagagagtgatcctggccgacgctaatctggacaaagtgctgtccgcctacaacaagcaccgggataagcccatcagagagcaggccgagaatatcatccacctgtttaccctgaccaatctgggagcccctgccgccttcaagtactttgacaccaccatcgaccggaagaggtacaccagcaccaaagaggtgctggacgccaccctgatccaccagagcatcaccggcctgtacgagacacggatcgacctgtctcagctgggaggcgacAAAAGGCCGGCGGCCACGAAAAAGGCCGGCCAGGCAAAAAAGAAAAAGtaa

#### SpCas9 variants, bacterial expression plasmids

Bacterial expression plasmids expressing SpCas9-HF1 and SuperFi-Cas9 were constructed from pET-Cas9-NLS-6xHis (Addgene #62933)^3^ plasmid in two steps. In the first step, the plasmid was digested with NcoI and PacI restriction enzymes and assembled with one PCR fragment using the NEBuilder HiFi DNA Assembly Master Mix. The PCR fragment was generated from pX330-Flag-WT_SpCas9 (without sgRNA; with silent mutations) (Addgene #126753) using the following primers: TCTAGAAATAATTTTGTTTAACTTTAAGAAGGAGATATACCATGGACAAGAAGTACAGCATCGGC and GGTGGCAGCAGCCTAGGTTATTAGTGATGGTGGTGGTGATGCTTATCCATCTTTTTCTTTTTTGCCTGGCCGG. SpCas9-HF1: In the second step, the first step product was digested with BglII and Eco52I restriction enzymes and assembled with one PCR fragment using the NEBuilder HiFi DNA Assembly Master Mix. The PCR fragment was generated from pX330-Flag-SpCas9-HF1 (without sgRNA; with silent mutations) (#126755) using the following primers:

GATACACCAGACGGAAGAACCGGATCTGCTATCTGCAAGA and CTTCTGCAGTTCGCCGGCAGAGGCC.

SuperFi-Cas9: In the second step, the first step product was digested with Eco72I and Mva1269I restriction enzymes and assembled with one PCR fragment using the NEBuilder HiFi DNA Assembly Master Mix. The PCR fragment was generated from pPIK16045_pX330_Flag-SuperFi-Cas9 (#184370) using the following primers:

GGAAACCCGGCAGATCACAAAGCACGTGGCACA and cagttcgccggcagaggc.

#### Other plasmid sequences

##### *Prnp*.HA-EGFP-DHFR[DD]

1,000 bp-long homology arm to the *Prnp* gene-EGFP-DHFR[DD] [EGFP-folA dihydrofolate reductase destabilization domain] fusion protein coding cassette-1,000 bp-long homology arm to the *Prnp* gene. Cloned from pBMN DHFR(DD)-YFP (#29325)^4^ using standard molecular biology techniques

Plasmid sequence:

acatgtgaagacaactacttgtgaatgcagctagatgtatctcagcagcccagcccgcccggctctcttgtctatggagagagtgaggaattctcaagtgagcctgccaaaactctagatgtttcctgtctctgataaacttaggttgaaaattccctcaaggagattcttggctttgtgcttaggggatgttaattccgtcacttgacagctgtgttgtgtcctcctctgtgccaggcactgcccttacccataaaatatggcaacgaaacagaggctcttggtttggtttggattctggggcatgagctgtaaagcccagatgtattagaactcacagccgtcctgtttcagcctctacttcccaagtgctggggcaccaacgtgcacttcctcatgcctggctctggagacctactgcttgtctccagggctcaaacactgagtcagctttcttcaagtccttgctcctgctgtagccactcaggagccctcctgactagaccatgactcaggcccttgtggtgttacggttactcaggaccagtgtactcacagctacccctgcaggtgactttctgcattctggggaatgaagcctacatccgtggataaaggttctcctctgtgtagaggctcacaccccacaggaccctggggccattatagcagccttatagtacagctgccaggctccccacaagatcatgcccatttccaaattccactacattgttaaagctcaaagccatggcgtaacaaccatgcaatatcacctagaccagacgtggtttaccagttggggtaactcttgtcaaatctgtcctcagaggatgggatgagctgtgtgtttttgatttacttttttcctgaaggaaaagctacggggggggggggggcgggggggagggttgacgccatgactttcatacatttgctttgtagatagatgtcaaggaccttcagcctaaatactgggcactgataccttgttcctcattttgcagatcagtcatcacacctgtcttcattaataccggtcgccaccatggtgagcaagggcgaggagctgttcaccggggtggtgcccatcctggtcgagctggacggcgacgtaaacggccacaagttcagcgtgtccggcgagggcgagggcgatgccacctacggcaagctgaccctgaagttcatctgcaccaccggcaagctgcccgtgccctggcccaccctcgtgaccaccctgacctacggcgtgcagtgcttcagccgctaccccgaccacatgaagcagcacgacttcttcaagtccgccatgcccgaaggctacgtccaggagcgcaccatcttcttcaaggacgacggcaactacaagacccgcgccgaggtgaagttcgagggcgacaccctggtgaaccgcatcgagctgaagggcatcgacttcaaggaggacggcaacatcctggggcacaagctggagtacaactacaacagccacaacgtctatatcatggccgacaagcagaagaacggcatcaaggtgaacttcaagatccgccacaacatcgaggacggcagcgtgcagctcgccgaccactaccagcagaacacccccatcggcgacggccccgtgctgctgcccgacaaccactacctgagcacccagtccgccctgagcaaagaccccaacgagaagcgcgatcacatggtcctgctggagttcgtgaccgccgccgggatcactctcggcatggacgagctGTACAagGGATCCCTtgcaGAGGCgTCtGCtAAAGCggcaGCgGGCTCGAGtATCAGTCTGATTGCGGCGTTAGCGGTAGATTACGTTATCGGCATGGAAAACGCCATGCCGTGGAACCTGCCTGCCGATCTCGCCTGGTTTAAACGCAACACCTTAAATAAACCCGTGATTATGGGCCGCCATACCTGGGAATCAATCGGTCGTCCGTTGCCAGGACGCAAAAATATTATCCTCAGCAGTCAACCGAGTACGGACGATCGCGTAACGTGGGTGAAGTCGGTGGATGAAGCCATCGCGGCGTGTGGTGACGTACCAGAAATCATGGTGATTGGCGGCGGTCGCGTTATTGAACAGTTCTTGCCAAAAGCGCAAAAACTGTATCTGACGCATATCGACGCAGAAGTGGAAGGCGACACCCATTTCCCGGATTACGAGCCGGATGACTGGGAATCGGTATTCAGCGAATTCCACGATGCTGATGCGCAGAACTCTCACAGCTATTGCTTTGAGATTCTGGAGCGGCGAtagCaattgttgttgttaacttgtttattgcagcttataatggttacaaataaagcaatagcatcacaaatttcacaaataaagcatttttttcactgcattctagttgtggtttgtccaaactcatcaatgtatcttaacgcgtcgtctctcctcgggaggccttcctgcttgttccttcgcattctcgtggtctaggctgggggaggggttatccacctgtagctctttcaattgaggtggttctcattcttgcttctctgtgtcccccataggctaatacccctggcactgatgggccctgggaaatgtacagtagaccagttgctctttgcttcaggtccctttgatggagtctgtcatcagccagtgctaacaccgggccaataagaatataacaccaaataactgctggctagttggggctttgttttggtctagtgaataaatactggtgtatcccctgacttgtacccagagtacaaggtgacagtgacacatgtaacttagcataggcaaagggttctacaaccaaagaagccactgtttggggatggcgccctggaaaacagcctcccacctgggatagctagagcatccacacgtggaattctttctttactaacaaacgatagctgattgaaggcaacaggaaaaaaaaaatcaaattgtcctactgacgttgaaagcaaacctttgttcattcccagggcactagaatgatctttagccttgcttggattgaactaggagatcttgactctgaggagagccagccctgtaaaaagcttggtcctcctgtgacgggagggatggttaaggtacaaaggctagaaacttgagtttcttcatttctgtctcacaattatcaaaagctagaattagcttctgccctatgtttctgtacttctatttgaactggataacagagagacaatctaaacattctcttaggctgcagataagagaagtaggctccattccaaagtgggaaagaaattctgctagcattgtttaaatcaggcaaaatttgttcctgaagttgctttttaccccagcagacataaactgcgatagcttcagcttgcactgtggattttctgtatagaatatataaaacataacttcaagcttatgtcttctttttaaaacatctgaagtatgggacgccctttctcgagacgcacaatgtgaaccatcaccctaatcaagttttttggggtcgaggtgccgtaaagcactaaatcggaaccctaaagggagcccccgatttagagcttgacggggaaagccggcgaacgtggcgagaaaggaagggaagaaagcgaaaggagcgggcgctagggcgctggcaagtgtagcggtcacgctgcgcgtaaccaccacacccgccgcgcttaatgcgccgctacagggcgcgtcaggtggcacttttcggggaaatgtgcgcggaacccctatttgtttatttttctaaatacattcaaatatgtatccgctcatgagacaataaccctgataaatgcttcaataatattgaaaaaggaagagtcctgaggcggaaagaaccagctgtggaatgtgtgtcagttagggtgtggaaagtccccaggctccccagcaggcagaagtatgcaaagcatgcatctcaattagtcagcaaccaggtgtggaaagtccccaggctccccagcaggcagaagtatgcaaagcatgcatctcaattagtcagcaaccatagtcccgcccctaactccgcccatcccgcccctaactccgcccagttccgcccattctccgccccatggctgactaattttttttatttatgcagaggccgaggccgcctcggcctctgagctattccagaagtagtgaggaggcttttttggaggcctaggcttttgcaaagatcgatcaagagacaggatgaggatcgtttcgcatgattgaacaagatggattgcacgcaggttctccggccgcttgggtggagaggctattcggctatgactgggcacaacagacaatcggctgctctgatgccgccgtgttccggctgtcagcgcaggggcgcccggttctttttgtcaagaccgacctgtccggtgccctgaatgaactgcaagacgaggcagcgcggctatcgtggctggccacgacgggcgttccttgcgcagctgtgctcgacgttgtcactgaagcgggaagggactggctgctattgggcgaagtgccggggcaggatctcctgtcatctcaccttgctcctgccgagaaagtatccatcatggctgatgcaatgcggcggctgcatacgcttgatccggctacctgcccattcgaccaccaagcgaaacatcgcatcgagcgagcacgtactcggatggaagccggtcttgtcgatcaggatgatctggacgaagagcatcaggggctcgcgccagccgaactgttcgccaggctcaaggcgagcatgcccgacggcgaggatctcgtcgtgacccatggcgatgcctgcttgccgaatatcatggtggaaaatggccgcttttctggattcatcgactgtggccggctgggtgtggcggaccgctatcaggacatagcgttggctacccgtgatattgctgaagagcttggcggcgaatgggctgaccgcttcctcgtgctttacggtatcgccgctcccgattcgcagcgcatcgccttctatcgccttcttgacgagttcttctgagcgggactctggggttcgaaatgaccgaccaagcgacgcccaacctgccatcacgagatttcgattccaccgccgccttctatgaaaggttgggcttcggaatcgttttccgggacgccggctggatgatcctccagcgcggggatctcatgctggagttcttcgcccaccctagggggaggctaactgaaacacggaaggagacaataccggaaggaacccgcgctatgacggcaataaaaagacagaataaaacgcacggtgttgggtcgtttgttcataaacgcggggttcggtcccagggctggcactctgtcgataccccaccgagaccccattggggccaatacgcccgcgtttcttccttttccccaccccaccccccaagttcgggtgaaggcccagggctcgcagccaacgtcggggcggcaggccctgccatagcctcaggttactcatatatactttagattgatttaaaacttcatttttaatttaaaaggatctaggtgaagatcctttttgataatctcatgaccaaaatcccttaacgtgagttttcgttccactgagcgtcagaccccgtagaaaagatcaaaggatcttcttgagatcctttttttctgcgcgtaatctgctgcttgcaaacaaaaaaaccaccgctaccagcggtggtttgtttgccggatcaagagctaccaactctttttccgaaggtaactggcttcagcagagcgcagataccaaatactgtccttctagtgtagccgtagttaggccaccacttcaagaactctgtagcaccgcctacatacctcgctctgctaatcctgttaccagtggctgctgccagtggcgataagtcgtgtcttaccgggttggactcaagacgatagttaccggataaggcgcagcggtcgggctgaacggggggttcgtgcacacagcccagcttggagcgaacgacctacaccgaactgagatacctacagcgtgagctatgagaaagcgccacgcttcccgaagggagaaaggcggacaggtatccggtaagcggcagggtcggaacaggagagcgcacgagggagcttccagggggaaacgcctggtatctttatagtcctgtcgggtttcgccacctctgacttgagcgtcgatttttgtgatgctcgtcaggggggcggagcctatggaaaaacgccagcaacgcggcctttttacggttcctggccttttgctggccttttgctc

##### *Sprn*.HA-CMV-EGFP

1,000 bp-long homology arm to the *Sprn* gene-CMV-EGFP protein coding cassette-1,000 bp-long homology arm to the Shadoo (*Sprn)* gene.

Plasmid sequence:

acatgtgaagacaactaacaaaatccagtcgtgagctctgcctaaagaaaagggtcctcgctgccgcacctttccgcttggccggtcaaggcccctaaatcgctatccgacctaggcttgtgaccagtaagtaggggaggtggatgaacccagtgatgctgggagggagggggaggggagagaggacctgaggaggatggagctgccgccaccgagatggctggtccacagccagccggaacccatcctgatcggtttacctgtccaggatactgccctggacaaacccaaaaagggtggagctggagcggggaagagacgtaattacctagggtctgggcctgctgggcggttcacccatcgctagttgtttgcgttgcaaggaatcgctgatctggattctgacccccaccctcacccagtgcattcagccgcagccactggctgaaagactatctcttaaggcatcaggagatccagatgccagagcaaaagtcacaaggctgtccattctaaccataatctgggggtattgaaggctctccattccaaactagaatcctaatccactaatccatccccttccagagactcgtgtgcagagcgggggatttgtgccccccccccaggcctgagccccactgtaggagctccgcaaaccccattctgggacccatctccaccctatcacaatagtaaaacggccgccctgtagttaaaccctttccccaccaccccatctccacctggttatacagcaggaacccaaaaccaaagtcctggtgctggagtttaaagaaccccttccccaaccctgagcccactgtcctccaagatgctgggagccctctatcggattaccacaaaaccagaggctgaagtaggtgtccccaggtccagatgagcctattcccaagccctgataccctcttgccctggtcctaaaccacgctccacccctgcacagaagctgaagccccttccaccctcttctcgcagattctgcccagtaggacacctgtcttcattaatagtaatcaattacggggtcattagttcatagcccatatatggagttccgcgttacataacttacggtaaatggcccgcctggctgaccgcccaacgacccccgcccattgacgtcaataatgacgtatgttcccatagtaacgccaatagggactttccattgacgtcaatgggtggagtatttacggtaaactgcccacttggcagtacatcaagtgtatcatatgccaagtacgccccctattgacgtcaatgacggtaaatggcccgcctggcattatgcccagtacatgaccttatgggactttcctacttggcagtacatctacgtattagtcatcgctattaccatggtgatgcggttttggcagtacatcaatgggcgtggatagcggtttgactcacggggatttccaagtctccaccccattgacgtcaatgggagtttgttttggcaccaaaatcaacgggactttccaaaatgtcgtaacaactccgccccattgacgcaaatgggcggtaggcgtgtacggtgggaggtctatataagcagagctggtttagtgaaccgtcagatccgctagcgctaccggtcgccaccatggtgagcaagggcgaggagctgttcaccggggtggtgcccatcctggtcgagctggacggcgacgtaaacggccacaagttcagcgtgtccggcgagggcgagggcgatgccacctacggcaagctgaccctgaagttcatctgcaccaccggcaagctgcccgtgccctggcccaccctcgtgaccaccctgacctacggcgtgcagtgcttcagccgctaccccgaccacatgaagcagcacgacttcttcaagtccgccatgcccgaaggctacgtccaggagcgcaccatcttcttcaaggacgacggcaactacaagacccgcgccgaggtgaagttcgagggcgacaccctggtgaaccgcatcgagctgaagggcatcgacttcaaggaggacggcaacatcctggggcacaagctggagtacaactacaacagccacaacgtctatatcatggccgacaagcagaagaacggcatcaaggtgaacttcaagatccgccacaacatcgaggacggcagcgtgcagctcgccgaccactaccagcagaacacccccatcggcgacggccccgtgctgctgcccgacaaccactacctgagcacccagtccgccctgagcaaagaccccaacgagaagcgcgatcacatggtcctgctggagttcgtgaccgccgccgggatcactctcggcatggacgagctgtacaagtccggactcagatctcgagctcaagcttcgaattctgcagtcgacggtaccgcgggcccgggatccaccggatctagataactgatcataatcagccataccacatttgtagaggttttacttgctttaaaaaacctcccacacctccccctgaacctgaaacataaaatgaatgcaattgttgttgttaacttgtttattgcagcttataatggttacaaataaagcaatagcatcacaaatttcacaaataaagcatttttttcactgcattctagttgtggtttgtccaaactcatcaatgtatcttaacgcgtcgtctctcctcaccaggctaaactccatcccaggtctagctcctagcctgtcttaaggcccctagggcccaccctaatggcctcctgcccagggggtaacttcattttgctaactatgattccccagcccagagaagagtctggccattctgggcccacaaggcctcttgcaatttgtggatgtagtttactctcttgcccatcccagtttcccagtcttccactctggcaggaactaacagccctactaccagagaggcaggctgcccttgactctccagaactgcccaagcaggcatgcctgtctgcttgcccaccctgacacgaggacatgaacaatgctgaccgccaagaagaggccatccttgggtgggcccgtctgattctgccactaaagcccccaaagaccttgagcctcccatggaaccagagggacttatagttgatgaagagagcattcacttagactgcagtgtgagaaatgtcaggtgctaaccacctgcccagagcaggtcccaatgatgtcatagctcagaaaacatccagagcagtaaaataaccatcccaggacgcccaccatgctcctcctacaacatgctaccaaagccaagtagtatgtttcttcctggcagactgcctaagagaccttgtgtcagaagggtttccacttgaagccacttggtcctaaaatccactgagggtagaggttttgaatacactttcaaaaacattccattctgcttgagcttaagggtcatgagtgagggtcacttggatataataccaatcctgctggggccttctttgtatataacccaaactgcaagattcccatagttccagtagatagcagcattttatgttgggagacccctcccttggaaacggttggacggggttggggtggggaggtgagacagagcatggcacagctgacagctggcaaactgaacagtggaaggggcagcagatctacagccccactgtgccagaacaaagctagcagacagattctcgagacgcactacgtgaaccatcaccctaatcaagttttttggggtcgaggtgccgtaaagcactaaatcggaaccctaaagggagcccccgatttagagcttgacggggaaagccggcgaacgtggcgagaaaggaagggaagaaagcgaaaggagcgggcgctagggcgctggcaagtgtagcggtcacgctgcgcgtaaccaccacacccgccgcgcttaatgcgccgctacagggcgcgtcaggtggcacttttcggggaaatgtgcgcggaacccctatttgtttatttttctaaatacattcaaatatgtatccgctcatgagacaataaccctgataaatgcttcaataatattgaaaaaggaagagtcctgaggcggaaagaaccagctgtggaatgtgtgtcagttagggtgtggaaagtccccaggctccccagcaggcagaagtatgcaaagcatgcatctcaattagtcagcaaccaggtgtggaaagtccccaggctccccagcaggcagaagtatgcaaagcatgcatctcaattagtcagcaaccatagtcccgcccctaactccgcccatcccgcccctaactccgcccagttccgcccattctccgccccatggctgactaattttttttatttatgcagaggccgaggccgcctcggcctctgagctattccagaagtagtgaggaggcttttttggaggcctaggcttttgcaaagatcgatcaagagacaggatgaggatcgtttcgcatgattgaacaagatggattgcacgcaggttctccggccgcttgggtggagaggctattcggctatgactgggcacaacagacaatcggctgctctgatgccgccgtgttccggctgtcagcgcaggggcgcccggttctttttgtcaagaccgacctgtccggtgccctgaatgaactgcaagacgaggcagcgcggctatcgtggctggccacgacgggcgttccttgcgcagctgtgctcgacgttgtcactgaagcgggaagggactggctgctattgggcgaagtgccggggcaggatctcctgtcatctcaccttgctcctgccgagaaagtatccatcatggctgatgcaatgcggcggctgcatacgcttgatccggctacctgcccattcgaccaccaagcgaaacatcgcatcgagcgagcacgtactcggatggaagccggtcttgtcgatcaggatgatctggacgaagagcatcaggggctcgcgccagccgaactgttcgccaggctcaaggcgagcatgcccgacggcgaggatctcgtcgtgacccatggcgatgcctgcttgccgaatatcatggtggaaaatggccgcttttctggattcatcgactgtggccggctgggtgtggcggaccgctatcaggacatagcgttggctacccgtgatattgctgaagagcttggcggcgaatgggctgaccgcttcctcgtgctttacggtatcgccgctcccgattcgcagcgcatcgccttctatcgccttcttgacgagttcttctgagcgggactctggggttcgaaatgaccgaccaagcgacgcccaacctgccatcacgagatttcgattccaccgccgccttctatgaaaggttgggcttcggaatcgttttccgggacgccggctggatgatcctccagcgcggggatctcatgctggagttcttcgcccaccctagggggaggctaactgaaacacggaaggagacaataccggaaggaacccgcgctatgacggcaataaaaagacagaataaaacgcacggtgttgggtcgtttgttcataaacgcggggttcggtcccagggctggcactctgtcgataccccaccgagaccccattggggccaatacgcccgcgtttcttccttttccccaccccaccccccaagttcgggtgaaggcccagggctcgcagccaacgtcggggcggcaggccctgccatagcctcaggttactcatatatactttagattgatttaaaacttcatttttaatttaaaaggatctaggtgaagatcctttttgataatctcatgaccaaaatcccttaacgtgagttttcgttccactgagcgtcagaccccgtagaaaagatcaaaggatcttcttgagatcctttttttctgcgcgtaatctgctgcttgcaaacaaaaaaaccaccgctaccagcggtggtttgtttgccggatcaagagctaccaactctttttccgaaggtaactggcttcagcagagcgcagataccaaatactgttcttctagtgtagccgtagttaggccaccacttcaagaactctgtagcaccgcctacatacctcgctctgctaatcctgttaccagtggctgctgccagtggcgataagtcgtgtcttaccgggttggactcaagacgatagttaccggataaggcgcagcggtcgggctgaacggggggttcgtgcacacagcccagcttggagcgaacgacctacaccgaactgagatacctacagcgtgagctatgagaaagcgccacgcttcccgaagggagaaaggcggacaggtatccggtaagcggcagggtcggaacaggagagcgcacgagggagcttccagggggaaacgcctggtatctttatagtcctgtcgggtttcgccacctctgacttgagcgtcgatttttgtgatgctcgtcaggggggcggagcctatggaaaaacgccagcaacgcggcctttttacggttcctggccttttgctggccttttgctc

1	Kulcsar, P. I. *et al.* Blackjack mutations improve the on-target activities of increased fidelity variants of SpCas9 with 5'G-extended sgRNAs. *Nat Commun* **11**, 1223, doi:10.1038/s41467-020-15021-5 (2020).

2	Engler, C., Kandzia, R. & Marillonnet, S. A One Pot, One Step, Precision Cloning Method with High Throughput Capability. *Plos One* **3**, doi:ARTN e3647

10.1371/journal.pone.0003647 (2008).

3	Zuris, J. A. *et al.* Cationic lipid-mediated delivery of proteins enables efficient protein-based genome editing in vitro and in vivo. *Nat Biotechnol* **33**, 73-80, doi:10.1038/nbt.3081 (2015).

4	Iwamoto, M., Bjorklund, T., Lundberg, C., Kirik, D. & Wandless, T. J. A general chemical method to regulate protein stability in the mammalian central nervous system. *Chem Biol* **17**, 981-988, doi:10.1016/j.chembiol.2010.07.009 (2010).
